# A shared cortical manifold links sensory error to motor planning

**DOI:** 10.64898/2026.08.02.742317

**Authors:** Grant W Zempolich, WenXi Zhou, Brooke E Holey, Alex H Williams, David M Schneider

**Affiliations:** Center for Neural Science, New York University, New York, NY 10012; Center for Computational Neuroscience, Flatiron Institute, New York, NY 10010

## Abstract

The ability to detect and correct errors is central to learning and executing skilled behaviors such as speech and musical performance. Accordingly, sensory and motor regions of the brain must communicate and coordinate their activity in order to both detect errors and adapt in response to them. However, the neural mechanisms by which sensory feedback is transformed into updated motor plans across distributed cortical circuits remain unclear. Here, we identify how the mouse brain encodes error signals during a skilled acoustic task and how these signals guide motor adaptation. We developed a novel, sound-dependent behavior in which mice use real-time auditory feedback to adjust ongoing forelimb movements. Task performance critically depends on auditory cortex, where neurons encode error-related feedback that predicts both within-trial and across-trial behavioral adaptations. Acoustic error feedback alters secondary motor cortex (M2) dynamics, selectively pushing activity along dimensions that encode planning signals for upcoming movements. Notably, auditory cortex and M2 share a low-dimensional manifold that emerges only after learning the skilled behavior. This shared geometry is not engaged during simpler forelimb tasks, suggesting that it reflects a learned, context-dependent solution for coordinating distributed cortical activity. In trained mice, silencing auditory cortex disrupts M2 dynamics, consistent with a model of continuously coupled cortical interactions during skilled performance. Together, these findings identify a learned coordinate transformation that maps sensory error onto corrective motor plans. More broadly, they point to a candidate general principle for how learning can establish a shared cortical manifold that links distant brain regions, enabling sensory feedback to selectively reshape future actions and support flexible, goal-directed control.

## Mice expertly perform a skilled acoustic behavior

Humans and other animals are adept at detecting acoustic errors and using them to guide rapid behavioral correction. During behaviors such as vocalization and musical performance, sensory feedback continuously refines motor output, allowing actions to be adjusted through trial and error ^1–3^. Yet, the cellular and circuit-level mechanisms by which the mammalian brain detects acoustic errors in real time and transforms them into specific changes in motor plans remain poorly understood.

Skilled behaviors such as musicianship highlight important interactions between hearing and motor centers of the brain during skilled acoustic behaviors. Musicians often describe a process of mental imagery that simulates upcoming gestures and anticipates their sounds, revealing a tight coupling between motor and auditory systems^4–6^. At the anatomical level, neural computations that support skilled acoustic behaviors are likely to involve the cerebral cortex, where auditory and motor centers are reciprocally connected ^1,7,89–12^. Neurophysiological studies show that during speech and manual sound production, the auditory cortex encodes sound, movement, and expectation, consistent with internal forward models that relate sounds to ongoing actions^13–18^. And when a behavior produces an unexpected sound, such as striking a wrong piano key, auditory cortex neurons generate mismatch responses that signal deviations from predicted outcomes^19–25^. These error-related signals can drive rapid corrective adjustments, implying that motor regions receive and use sensory feedback to adapt behavior. Indeed, current evidence now positions sensation as an important contributor to motor dynamics^2,12,26–29^.

Although prior findings point to a distributed cortical network that integrates sensory, motor, and predictive information to support skilled acoustic behavior, how the brain detects acoustic errors and converts them into adaptive motor commands remains unknown. To begin investigating the neural circuits underlying skilled acoustic behavior, we trained mice to perform precise forelimb movements guided by closed-loop auditory feedback (Fig. 1a,b). Head-restrained mice were tasked with pushing a lever from a position near their body (home) forward into a narrow target zone (1.77mm wide). Mice heard a single tone when the lever entered the target zone (entry tone, 16kHz), and a second tone if the lever exceeded the target zone (exit tone, 8kHz). Lever presses that peaked within the target zone (correct trials) produced only the entry tone and yielded a small water reward when the lever was returned to the home position. Lever presses that peaked too short (undershoots) produced no tones and lever presses that peaked too long (overshoots) produced both the entry and exit tones; neither undershoot nor overshoot trials were rewarded. Every 30 trials, the target zone was moved to 1 of 3 non-overlapping locations (chosen pseudorandomly), requiring mice to regularly update their behavior using acoustic feedback (Fig. 1c; Extended Data Fig. 1a).

**Figure 1:**
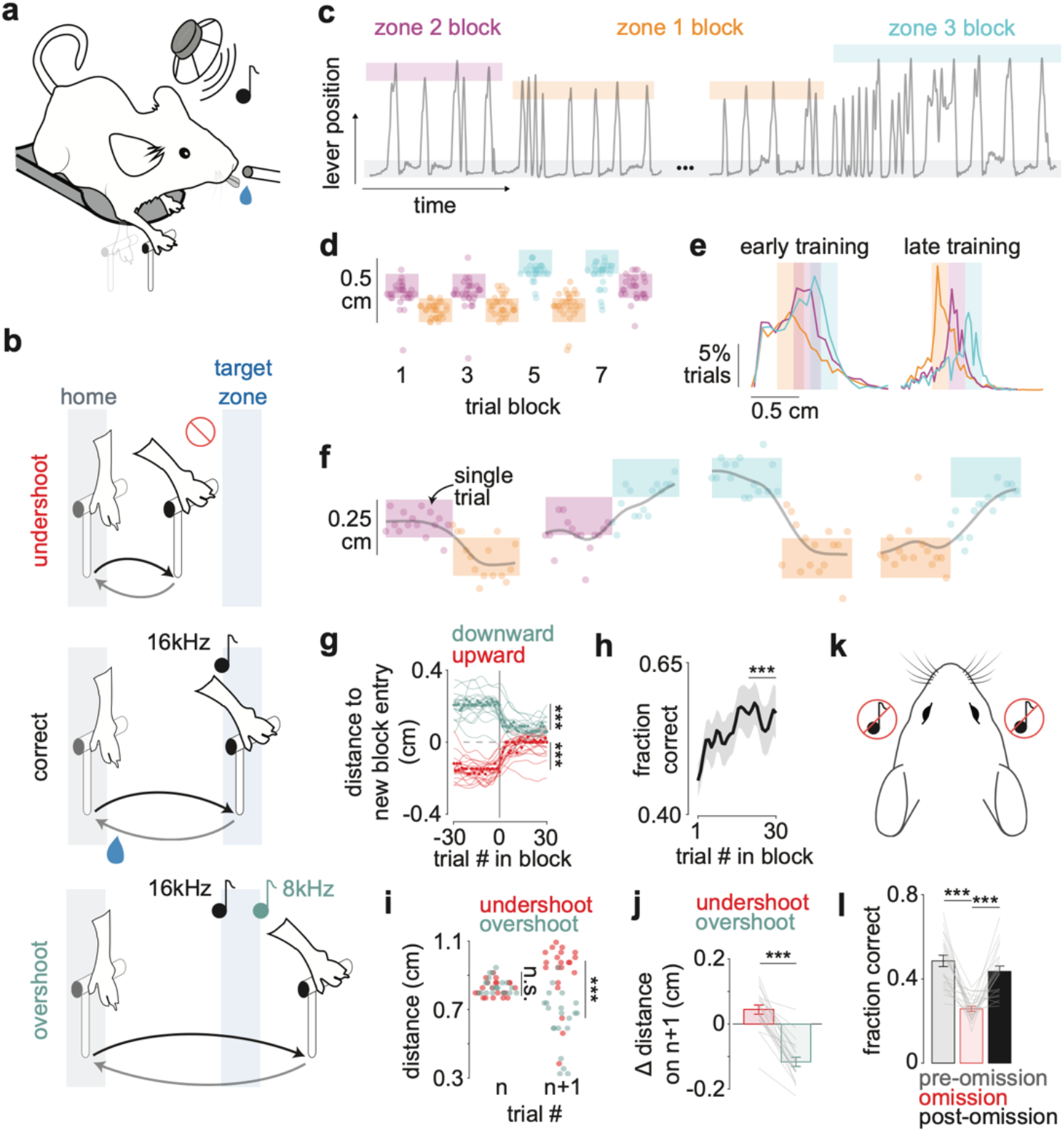
Mice perform a skilled acoustic forelimb behavior. **a**, Schematic depicting behavioral paradigm design. **b**, Schematic depicting closed-loop rules governing the task. Only trials that elicit the single entry tone (16kHz) are correct and rewarded. **c**, Example behavioral data showing lever position over time. Gray rectangle shows the home position; colored rectangles show the target zone on different blocks of trials. **d**, Example behavioral data with individual trials represented as dots (dots depict the peak of a each press). Shaded rectangles represent the target zones, and dots that land within the corresponding target zone are correct trials. **e**, Peak press distributions early and late in training across mice (N = 20). Colors show data sorted by target zone location. **f**, Example behavior aligned to zone transitions (15 trials before and after each transition). Gray lines depict moving averages of lever peak across trials. **g**, Population average data showing change in peak lever press around near-to-far (“upward”) and far-to-near (“downward”) zone transitions (N = 20 mice; ranksum, ***p<0.001). Colored lines represent individual mice; dots represent population average. **h**, Mean performance across the 30 trials within a block (N = 20 mice). Performance on the last 10 trials is significantly higher than the first 10 trials within a block (ranksum, ***p<0.001). Shaded regions represent standard error of the mean (SEM). **i**, Example data from a single mouse showing press amplitudes on errors that ended in Zone 2 range (trial n) and on the subsequent trial (trial n+1) (ranksum, ***p<0.001; n.s. not-significant). **j**, Change in peak amplitude on trials following undershoot and overshoot errors; data shown for all mice (N=20, paired t-test, ***p<0.001). **k**, Schematic depicting tone omission experiments. **l**, Fraction correct during omission blocks and the control blocks surrounding the omission blocks, (N=20, paired t-test, ***p<0.001). Error bars represent SEM.

Across 2 to 3 weeks of practice, mice learn to reliably press the lever into the target zone and to adapt their behavior following zone transitions (Fig. 1d). Performance improves with training despite the target zones becoming progressively smaller and non-overlapping across days (2.80+/-0.38mm in early training; 1.77+/-0mm at the end of training), and mouse performance reaches behavioral levels significantly above chance performance in all three zones (Fig. 1e; Extended Data Fig. 1b,c). Once mice reach expert-level performance, they continue to rely on acoustic feedback to adjust their behavior following errors induced by unexpected target zone transitions (Fig. 1f). When the zone location shifts nearer or farther away, mice adjust their pressing amplitude accordingly, producing changes that mirror the direction of the shift and lead to improved performance over the course of the block. After an overshoot trial, mice tend to reduce their press amplitude on the subsequent attempt, whereas after an undershoot, they increase it (Fig. 1g,h). These across-trial corrections cannot be accounted for by simple regression to the mean press amplitude across a session, since undershoots and overshoots within the range of zone 2 (that is, with equivalent amplitudes) elicit corrective adjustments in opposite directions on the subsequent trial (Fig. 1i,j). Furthermore, mice gradually learn to decelerate just prior to reaching the target zone, demonstrating an ability to anticipate the zone’s location through experience (Extended Data Fig. 2a-d). These directed behavioral adjustments demonstrate that expert mice flexibly update their actions in response to specific acoustic feedback cues to optimize their performance.

**Figure 2:**
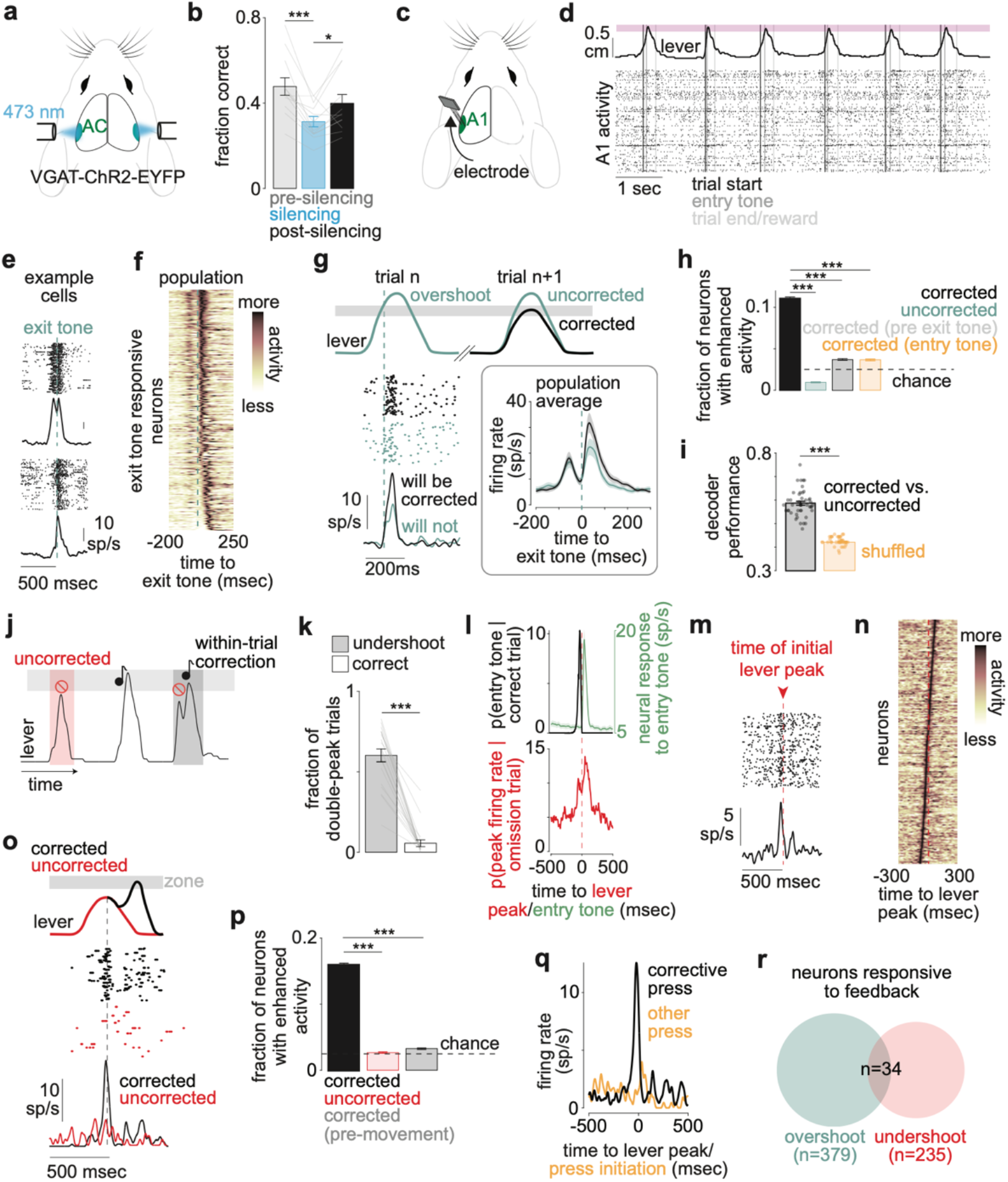
Neural activity in auditory cortex signals behavioral adaptations. **a**, Schematic depicting bilateral auditory cortex optogenetic silencing experiments. **b**, Fraction correct shown for VGAT-ChR2-EYFP mice during optogenetic perturbation blocks and the control blocks surrounding the perturbation blocks, (N=12, paired t-test, ***p<0.001, *p<0.05). Error bars represent SEM. **c**, Schematic depicting A1-targeted electrophysiology in head-fixed mice. **d**, Example experimental data showing population electrophysiology (bottom) and lever position (top). Colored rectangle is the target location. **e**, Example neurons responsive to the exit tone on overshoot trials. **f**, Trial averaged PSTHs for A1 neurons exhibiting significant responses to the exit tone (0ms to 100ms) on overshoot trials (n=379, rank-sum compared to baseline −300ms to −200ms, p<0.05). Each neuron’s PSTH normalized to occupy a range of 0-1. **g**, Top: schematic depicting division of overshoot trials that result in future correction versus those that remain uncorrected. Bottom left: example neuron showing an increased response to the exit tone on trials preceding behavioral correction. Bottom right: average population response of exit tone responsive neurons favoring correction (n=45) (see Methods). Shaded regions represent SEM. **h**, Two-sided permutation test (see Methods) identifying percent of neurons preferring corrected (black) versus non-corrected (green) conditions using neural activity 0ms to 50ms after exit tone onset. “Corrected pre-exit tone” (gray) indicates equivalent analysis using neural activity −50ms to 0ms preceding exit tone onset. “Corrected entry tone” (orange) indicates equivalent analysis using neural activity 0ms to 50ms surrounding entry tone onset during the same overshoot trials. Neurons with enhanced activity were identified with a two-sided permutation test with an alpha value of 0.05. (ranksum, ***p<0.001). Error bars represent SEM. **(continued)**. **l**, Top: Histogram showing probability distribution of entry tone times relative to lever apex (time zero) on correct trials (black); A1 population neural response to entry tone for neurons exhibiting significant increases in spiking (0ms to 100ms) on correct trials (green) (n=414, rank-sum compared to baseline −300ms to −200ms, p<0.05). Bottom: probability distribution of neurons displaying significant increase in firing rate at each time point −500ms to 500ms surrounding undershoot apex. 50ms sliding window advanced in 10ms increments (see Methods). **m**, Example neuron responsive to the omission of the entry tone aligned to the initial apex on undershoot trials regardless of future correction. **n**, Trial averaged PSTHs of A1 neurons exhibiting peak responses between −75ms to +75ms surrounding the apex of undershoot trials (n=235). Each neuron’s PSTH normalized to occupy a range of 0-1. **o**, Example neuron showing an increased response on undershoot trials with a subsequent within-trial correction. **p**, Percent of neurons that respond more strongly on trials that will be corrected (black) versus those that will not be corrected (red). Activity before movement is not modulated by whether or not a trial will be corrected (gray). Dashed line represents 2.5% chance level (ranksum, ***p<0.001). Error bars represent SEM. **q**, Trial-averaged PSTHs for an example neuron, aligned to undershoot apexes during within-trial corrected movements (black) versus trial initiation of correct trials (i.e. non-undershoot event trials) in orange. **r**, Venn diagram showing A1 neurons that encode overshoot (exit tone) and undershoot (tone omission) events.

Although our task design was performed in darkness and provided explicit acoustic cues to guide behavior, it remains possible that mice adopt strategies that rely on other cues (e.g. proprioceptive feedback). To test this, we omitted sounds during the entirety of select blocks of trials (every 1 of 3 blocks) in expert mice and observed their behavior (Fig. 1k; Extended Data Fig. 3a). During omission blocks, mice still received rewards for apexing the lever within the target zone, but they received no acoustic feedback. With sounds omitted, performance dropped toward chance levels, and behavior recovered immediately during the next sound-generating block (Fig. 1l; Extended Data Fig. 3b,c). Although mice sometimes apexed their lever movement within the target zone during omission trials and received a reward, they failed to maintain this behavior on subsequent trials, indicating that mice rely upon acoustic cues to maintain correct behavior (Extended Data Fig. 3d). Altogether, these data show that mice use real-time auditory feedback to guide their actions and learn from mistakes during a skilled acoustic behavior.

**Figure 3:**
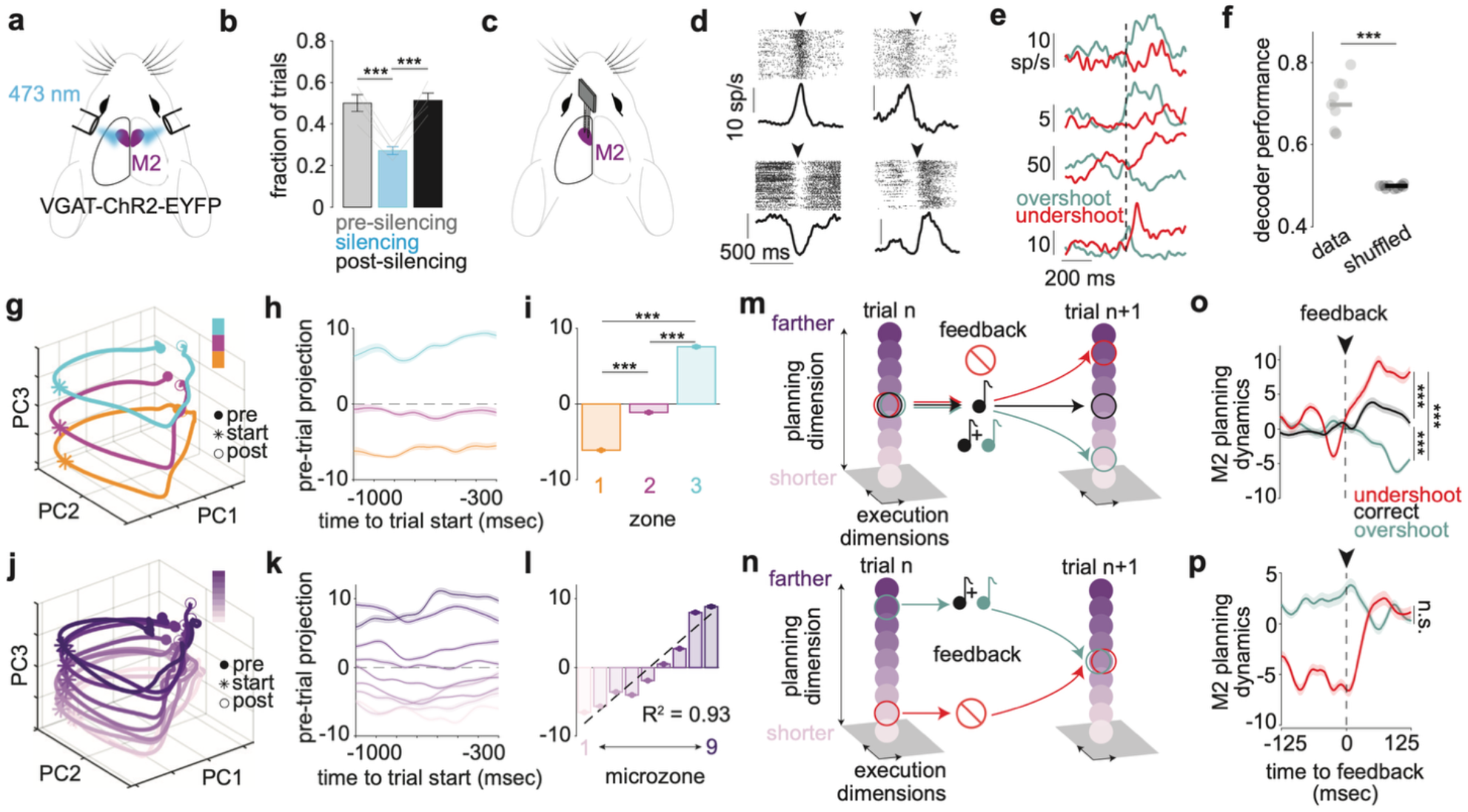
Acoustic feedback impacts M2 population dynamics for motor planning. **a**, Schematic depicting bilateral secondary motor cortex (M2) optogenetic silencing. **b**, Average performance for VGAT-ChR2-EYFP mice during optogenetic perturbation blocks and the control blocks surrounding the perturbation blocks, (N=4, paired t-test, ***p<0.001). Error bars represent SEM. **c**, Schematic depicting M2-targeted electrophysiology in head-fixed mice. **d**, Example M2 neurons aligned to trial initiation (black arrow). **e**, Example M2 neurons with sustained activity following either overshoot (top 2 neurons) or undershoot (bottom 2 neurons) errors (ranksum test, p<0.05). **f**, Decoder performance (predicting error type) trained on neural activity from individual recordings during the post-error interval preceding the next trial (N=9, ranksum ***p<0.001). Horizontal bars represent population means. **g**, Population-level neural data projected onto top 3 principal components (conditioned by target zone). Trajectories show 500 ms preceding movement onset (pre) through 1000 ms following movement onset (post). **h**, Population-level neural data during pre-trial intervals (in the absence of movement) projected onto the top principal component derived from pre-trial activity. Shaded regions represent 95% CI across test/train splits (see Methods). **i**, Quantification of projection values (a.u.) over full analysis window shown in (h) (ranksum, ***p<0.001). **j**, Population-level neural data projected onto top 3 principal components (conditioned by microzone). Data show 500 ms preceding movement onset (pre) through 1000 ms following movement onset (post). **k**, Population-level neural data during pre-trial intervals (in the absence of movement) projected onto the top principal component derived from pre-trial activity, for microzone-sorted data. Shaded regions represent 95% CI across test/train splits (see Methods). **l**, Linear regression on mean projection values over analysis window shown in (k) (R2 =0.93, p<0.001). **m**, Schematic depicting model in which time points of feedback reset the initial conditions away from the intermediate pressing range for subsequent movements along a motor planning dimension of M2 neural activity. **n**, Schematic depicting a model in which time points of feedback reset the initial conditions towards the intermediate pressing range for subsequent movements along a motor planning dimension of M2 neural activity. **o**, M2 activity (a.u.) projected onto the planning axis (PC1 of pre-trial neural activity), aligned to the moment of feedback during correct (time of entry tone), undershoot (omission of tone at initial apex), and overshoot (time of exit tone) trials. All data are from lever presses that apexed within Zone 2 range, sorted by trial outcome. Shaded regions represent 95% confidence interval across bootstraps (see Methods), (ranksum ***p<0.001). **p**, As in (**o**), but for undershoot and overshoot trials that were intended to land in Zone 2 (ranksum, n.s. not-significant). Correct trial curve not shown; data is equivalent to the data previously shown in (**o**).

## Auditory cortex activity predicts overshoot behavioral adaptations

The auditory cortex integrates sound and movement-related signals and is an important locus for forming internal models relating action to acoustic outcome, making it a potentially important brain area for skilled acoustic behaviors like the one described above ^18,30,31^. However, acoustic tasks involving tones can often be performed without auditory cortex, and it remains unknown whether auditory cortex is necessary for behaviors that are guided by simple acoustic feedback yet still require skill ^32,33^. To test whether the auditory cortex is necessary for this skilled acoustic task, we performed behavior-triggered, bilateral silencing of auditory cortex during a subset of trial blocks (Fig. 2a; Extended Data Fig. 4a). During these perturbation blocks, mouse-initiated lever presses triggered the closed-loop activation of a 473nm laser for the duration of each lever press, thus stimulating auditory cortex inhibitory cells and suppressing activity in local excitatory cells (in VGAT-ChR2-EYFP mice). In VGAT+ mice but not in WT/VGAT-controls, performance decreased toward chance levels with auditory cortex silenced, and mice were unable to adapt their behavior to match the zone location, revealing the necessity of auditory cortex for detecting acoustic error feedback and learning from mistakes (Fig. 2b; Extended Data Fig. 4b-f).

**Figure 4:**
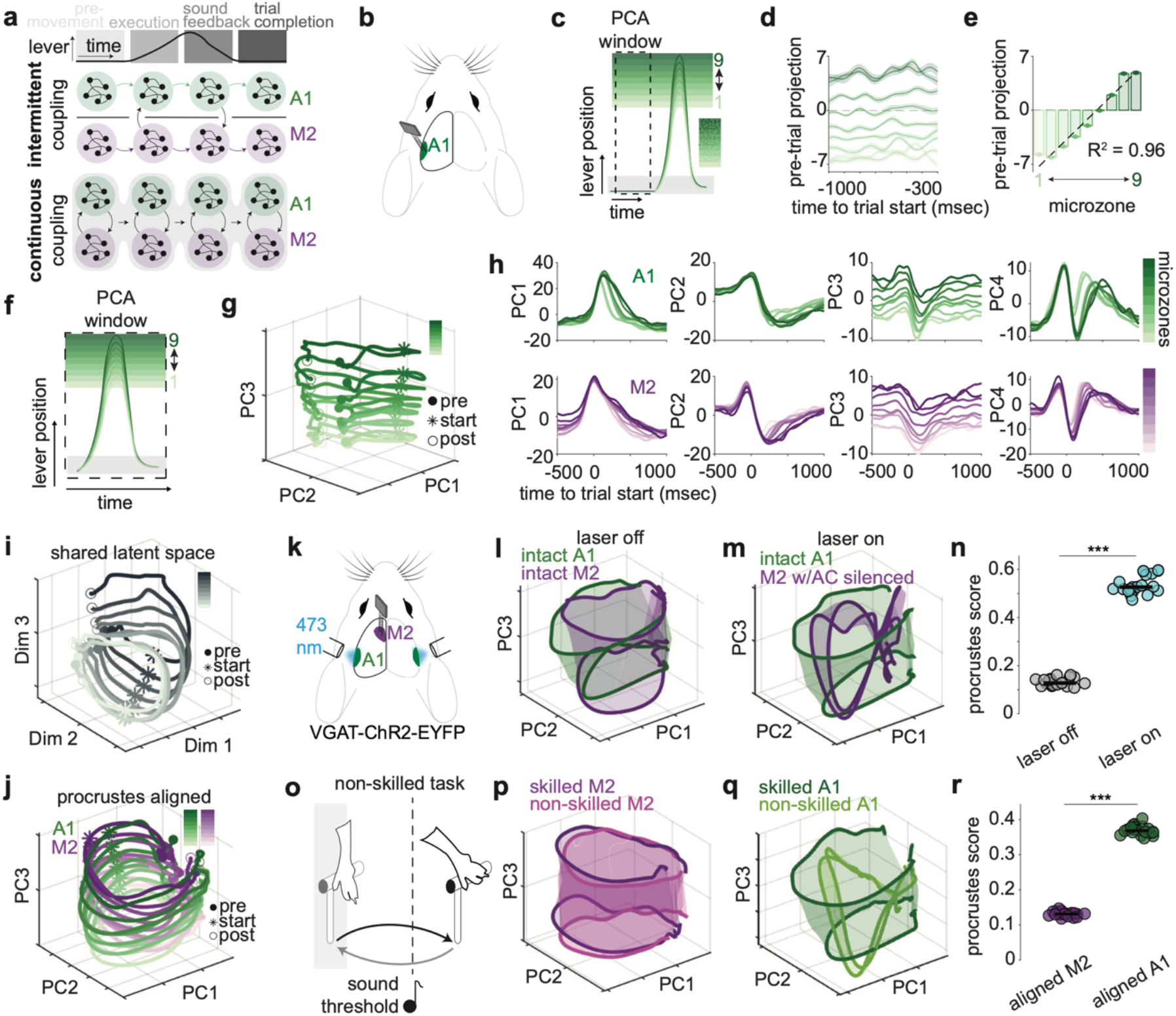
Auditory cortex and M2 share experience-dependent manifold geometries. **a**, Top: Schematized single trial press during skilled, auditory-guided behavior. Bottom: Schematics depicting opposing models of coupling between M2 and A1 during behavior. **b**, Schematic depicting A1-targeted electrophysiology in head-fixed mice. **c**, Schematic depicting activity window used for PCA training and results depicted in (**d**) and (**e**). **d**, Projections of microzone-specific neural activity (a.u.) onto top principal component derived from pre-trial activity. Shaded regions represent 95% CI across test/train splits (see Methods). **e**, Linear regression on mean projection values (a.u.) over analysis window shown in (**d**) (R2 =0.96, p<0.001). **f**, Schematic depicting activity window used for PCA training and results depicted in (**g**). **g**, Projection (mean across test/train splits) of nine-microzone split neural data into subspace defined by top 3 principal components derived from trial-start aligned neural activity. Pre (−500 ms), post (+1000 ms). **h**, Low-dimensional projections (mean across test/train splits) of A1 (n=1164) and M2 (n=632) population activity (a.u.) onto top 4 principal components computed separately for each brain region. A1, 27.89% variance explained by top 4 PCs; M2, 33.7% variance explained by top 4 PCs. Shaded regions represent 95% CI across test/train splits. **i**, Mean latent space projection into A1-M2 joint subspace identified via CCA. Projections for both regions averaged together and shown as individual condition trajectories. Pre (−500 ms), post (+1000 ms). **j**, Procrustes aligned 3D neural manifolds (0.18 score +/-0.001), A1 (green) and M2 (purple). Reference region: M2. **k**, Schematic depicting bilateral silencing of AC while recording in M2 during behavior in VGAT(+) mice. **l**, Procrustes aligned 3D neural manifolds (0.13 score +/-0.014), intact A1 (green) vs. intact M2 (purple). Reference region: intact A1. Manifolds interpolated across amplitude conditions to assist with visualization. **m**, Procrustes aligned 3D neural manifolds (0.53 score +/-0.041), intact A1 (green) vs. M2 with AC silenced (purple). Reference region: intact A1. **n**, Procrustes alignment quantification across 20 data splits, comparing laser off/on conditions, (rank sum, ***p<0.001). Each dot represents alignment metric of neural manifolds in the top 3 PC subspace trained on a single data split (see Methods). Horizontal bars represent population means. **o**, Schematic depicting simple, non-skilled lever pressing task. **(continued)**. **p**, Procrustes aligned 3D neural manifolds (0.13 score +/-0.006), skilled M2 (n=632) vs. non-skilled M2 (n=1326) neural populations. Reference region: skilled M2. **q**, Procrustes aligned 3D neural manifolds (0.36 score +/-0.019), skilled A1 (n=1164) vs. non-skilled A1 (n=409) neural populations. Reference region: skilled A1. **r**, Procrustes alignment quantification across 20 data splits, comparing M2 (skilled vs. non-skilled) and A1 (skilled vs. non-skilled), (rank sum, ***p<0.001). Each dot represents alignment metric of neural manifolds in the top 3 PC subspace trained on a single data split (see Methods). Horizontal bars represent population means.

To investigate how primary auditory cortex (A1) neurons encode acoustic feedback related to error, we performed electrophysiological recordings in expert mice performing the task (Fig. 2c,d; Extended Data Fig. 5a). We first focused on overshoot trials, since these trials provide a distinct moment of acoustic feedback (the time of the exit tone). Aligning neural activity to the exit tone on overshoot trials, we found that 28.2% of A1 neurons respond significantly above baseline to the exit tone (n=379, rank-sum, p<0.05) (Fig. 2e,f).

One important test of whether responses to the exit tone represent error is whether the activity of these exit-tone-responsive neurons correlates with changes in behavior. Specifically, we wondered whether A1 responses to error-related acoustic feedback on overshoot trials predict whether or not a mouse will make a behavioral adaptation on the following trial. To assess this, we separated overshoot trials (trial n) into two groups based on whether mice fix their behavior on the next trial (trial n+1) or whether they continued to overshoot. Importantly, both groups of initial overshoot trials were behaviorally and acoustically equivalent, with the only difference being what mice did on the next trial (~0.4 to 1 seconds in the future). Exit-tone responsive neurons were significantly more responsive to the exit-tone on overshoot trials that preceded a behavioral adaptation compared to overshoot trials that were not subsequently corrected (Fig. 2g,h). Using the trial-by-trial responses to the exit tone of individual neurons, we could decode whether a mouse would adapt its behavior significantly above chance (Fig. 2i). Neural activity during an early phase before exit-tone playback failed to predict these across-trial changes in behavior, nor did neural responses aligned to the entry tone, indicating that neural activity predictive of behavioral changes is not due to a state-like change in responsiveness on some trials compared to others (e.g. due to attention or arousal) (Fig. 2h; Extended Data Fig. 5b).

## Auditory cortex activity predicts undershoot behavioral adaptations

In addition to adapting their behavior across trials, mice sometimes make within-trial corrections immediately following undershoots, similar to the rapid behavioral corrections performed by skilled musicians ^34,35^ (Fig. 2j). Our task design allowed mice to salvage an undershoot trial by pushing the lever back toward the target zone, so long as they did so before returning the lever to the home position. On 58% of undershoot trials (range: 17-83%), mice turned the lever around within 70+/-1ms of their original apex, and a majority of these turn-around trials (57%; range: 21-87%) entered the target zone and led to reward, termed “within-trial corrected” presses. Such multi-peaked movements were rarely observed on correct trials following the zone entry apex, indicating that mice are not generally prone to making “double-takes” after they make an initial apex, but do so as a reaction to the absence of acoustic feedback on undershoot trials (Fig. 2k). Neural responses during undershoot trials were not distributed evenly throughout the movement, but were concentrated during a narrow window surrounding the initial apex (−75ms to +75ms) that coincided with when the tone would have been heard on a correct trial, consistent with these neurons signaling the omission of an expected acoustic outcome (Fig. 2l). Within this window, we identified a substantial population of A1 neurons that exhibited enhanced activity around the initial peak of the mouse’s lever press on undershoot trials (n=235) (Fig. 2m,n). To determine whether A1 activity can predict rapid, within-trial behavioral corrections following undershoots, we sorted these undershoot trials into those that were corrected within-trial versus those that were uncorrected. Omission-encoding neurons exhibited significantly elevated activity at the time of sound omission on within-trial corrected trials relative to uncorrected trials (Fig. 2o,p). As observed during overshoot corrections, strong activity during within-trial corrected omission trials does not reflect a global change in neuronal excitability, but instead is specific for the moment of error (Fig. 2p; Extended Data Fig. 5c). In addition to predicting rapid within-trial changes in behavior, omission-related activity in A1 also encodes across-trial changes in behavior, with larger responses at the moment of omitted feedback during uncorrected undershoot trials that precede behavioral corrections on the next trial (Extended Data Fig. 5d). Omission-encoding neurons were not active during windows preceding non-corrective lever movements (i.e. trial initiation on correct presses), indicating that their activity is dissociable from motor execution (Fig. 2q; Extended Data Fig. 5e).

Notably, auditory cortex neurons encoding overshoot and undershoot errors are almost completely non-overlapping, indicating that distinct populations of neurons in mouse A1 encode distinct types of error-related feedback during skilled acoustic behavior (Fig. 2r). These data are consistent with auditory cortex encoding error-related acoustic feedback that is used to guide behavior.

## Secondary motor cortex encodes acoustic errors

Although the auditory cortex is required for skilled acoustic behavior and responds to acoustic error feedback, it likely contributes to skilled behavior via its connections with motor systems in the brain ^10,12,36^. The motor cortex is important for many sensory-driven behaviors ^37–40^. In mice, the secondary motor cortex (M2) is reciprocally connected with A1, and A1 neurons send sound-related signals to M2, suggesting that M2 may play a role in using acoustic feedback to adaptively update behavior ^10,12^.

To test whether M2 activity is needed during this skilled acoustic behavior, we performed behavior-triggered optogenetic silencing of bilateral M2 (Fig. 3a; Extended Data Fig. 6a). We found that M2 was critical for the successful execution of the task, consistent with M2 playing an important role in skilled, sensory-guided behavior (Fig. 3b; Extended Data Fig. 6b-f). We next made electrophysiology recordings in M2 of expert mice during behavior (Fig. 3c). While population activity peaked around trial onset, individual M2 neurons exhibited diverse task-related modulation patterns, including both increases and decreases in activity. These distinct profiles suggest heterogeneous encoding and population-level task representations in M2 during behavior (Fig. 3d; Extended Data Fig. 7a).

Many M2 neurons (n=145, rank-sum, p<0.05) have rapid, short-latency responses to passive tones presented outside of the context of behavior, reminiscent of tone-evoked responses observed in A1 and consistent with the routing of sound-related information to motor cortex. However, these transient sound-evoked responses in M2 are almost completely gated off during behavior, consistent with previous reports (Extended Data Fig. 7b,c) ^12^. In contrast to the transient acoustic feedback signaling observed in auditory cortex during behavior, activity in many M2 neurons (n=158, rank-sum test, p<0.05) displays a more sustained increase following errors, with individual neurons preferring different error types (Fig. 3e). Across the M2 population, error type could be decoded significantly above chance based on neural activity during the inter-trial interval following an error (Fig. 3f). Differences in firing rate between error conditions cannot be accounted for by differences in other task-related behaviors such as licking (Extended Data Fig. 7d). These data reveal that M2 activity is impacted by acoustic information during behavior and that unlike the transient error encoding observed in A1, acoustic errors drive more persistent signaling in M2 that distinguishes between error types.

## Acoustic error feedback selectively impacts M2 dynamics for motor planning

Having established that acoustic errors elicit persistent modulation of M2 activity, we next examined how these signals interact with motor planning and execution. We first applied principal component analysis (PCA) to trial-averaged neural activity pooled across subjects (N=9, n=676) and aligned to lever presses to each of the three target zones (Fig. 3g; Extended Data Fig. 7e). This revealed that distinct neural trajectories for each target were offset from one another within 3-dimensional PC space (47% variance explained, top 3 PCs), not only during movement execution but also during the period prior to movement onset (Fig. 3h,i; Extended Data Fig. 7f). Does this population activity reflect abstract categorical representations of the zone a mouse plans to target with the lever, or does it reflect more fine-grained aspects of movement planning? To address this question, we further subdivided trials into nine groups of different lever-press magnitudes by partitioning each of the three original target zones into three equally sized microzones (Extended Data Fig. 7g). Applying the same PCA analysis to the nine-microzone data revealed a finely graded, linear organization of neural activity as a function of lever-press magnitude (29.4% variance explained, top 3 PCs; Fig. 3j-l), indicating that M2 dynamics encode fine-grained motor planning rather than coarse target categories. Notably, trajectories for different amplitude presses were systematically displaced along an orthogonal dimension prior to and during movement, closely paralleling population dynamics previously described in motor cortex during a variety of behaviors ^41^. These findings are consistent with a model in which rotational dynamics in M2 are correlated with ongoing motor control and that translation along a third, putative output-null dimension represents planning for different movement variations (i.e. lever press amplitude) ^37,41–45^.

Given that mice use errors to guide future behavior during this task, we hypothesized that acoustic feedback should modulate M2 dynamics related to motor planning. In particular, we reasoned that timepoints of error feedback should modulate M2 dynamics by selectively moving activity along the planning-related dimension, thus resetting the initial conditions for subsequent movements (Fig. 3m,n).

To investigate whether error feedback interacts with this planning-related dimension, we first analyzed trials in which lever presses terminated in the range of zone 2, regardless of the intended target. This allowed us to compare neural responses across equivalent movements that nonetheless produced different types of feedback: correct trials (intended for zone 2) produced only the entry tone, overshoots (intended for zone 1) produced the entry and exit tones, and undershoots (intended for zone 3) resulted in sound omission. Projection of M2 population activity onto the principal component that encodes movement planning revealed that timepoints of feedback induced large, directionally consistent deflections along this axis, without impacting execution-related dimensions of neural activity (Fig. 3o; Extended Data Fig. 8a-h; Extended Data Fig. 9a; Extended Data Fig. 10a,c,e,g). Following the exit tone on overshoot errors, activity was rapidly displaced toward the neural state associated with shorter press amplitudes. Conversely, the timepoint of initial apex during undershoot trials (i.e., omission of a tone) induced a rapid shift toward neural states corresponding to longer presses. Hearing only the entry tone on correct trials had a negligible effect on motor planning dynamics. When examining trials in which animals incorrectly landed in the range of zones 1 or 3 but had been targeting zone 2, we observed the reverse pattern: error-aligned deflections shifted M2 planning dynamics back toward intermediate sized movements, consistent with behavioral updating in the opposite direction (Fig. 3p). As before, execution-related dimensions of neural activity were not impacted by acoustic feedback and displayed equivalent dynamics for the course of the entire trial (Extended Data Fig. 8i-l; Extended Data Fig. 10b,d,f,h).

Together, these results indicate that acoustic error signals selectively modulate a motor planning dimension in M2, consistent with the implementation of a rapid, learned coordinate transformation that links acoustic feedback to adaptive updating of movement planning.

## Auditory cortex and M2 share experience-dependent manifold geometries

In musicians, experience with an instrument leads to strong coupling of motor and auditory circuits in the brain ^6,46^. Experimental and computational work suggests that continuous coupling between motor and sensory regions can facilitate communication through the emergence of shared manifold geometries, acting as a substrate for computations that require cross-modal integration, and may arise only when such cross-modal integration is necessary (e.g. as in trained musicians) ^47–51^. In contrast, traditional models of sensorimotor behaviors posit a more intermittent coupling, with sensory and motor regions operating largely autonomously, but transmitting information between one another at discrete moments (e.g. moments of behavioral execution, sensory feedback, or reinforcement) ^21,52,53^ (Fig. 4a).

A continuous-coupling model of auditory-motor computation yields four testable predictions: A1 and M2 dynamics should remain correlated continuously throughout behavior, not just during discrete events (e.g. moments of acoustic feedback); if coupling aligns population activity across areas, A1 and M2 population dynamics might exhibit similar manifold geometry; silencing one area should disrupt dynamics in the other; and shared dynamics should be prominent only in animals that have learned a behavior that would benefit from aligned manifold geometry.

We first quantified the structure of task-related activity in A1 by applying dimensionality reduction to population activity during a window preceding movement onset (Fig. 4b,c). Doing so revealed a finely graded, linear organization of neural activity as a function of future lever-press magnitude, reminiscent of motor planning signals observed in M2 (Fig. 4d,e). Extending the analysis window to include neural activity during presses to each microzone (−500 ms to +1000 ms surrounding trial initiation) revealed a tubular manifold embedded in A1 activity during behavior, bearing striking similarity to that observed in M2 (Fig. 4f,g; Extended Data Fig. 11a,b).

We next visualized side by side the low-dimensional population activity in A1 and M2 during behavior. Doing so revealed a strong correspondence between latent projections in each area that is maintained prior to and throughout the duration of behavior. Although there were small temporal offsets between the population dynamics in each area, the top principal components of activity evolved in parallel with one another prior to and throughout behavior, consistent with coupled dynamics (Fig. 4h). As observed in M2, error-related feedback also modulated motor-planning-related activity in A1. These deflections emerged with a delay of approximately 80-100 milliseconds, consistent with updated motor plans arising first in M2 and then being routed to A1 (Extended Data Fig. 12a,b). Together, these findings support the notion that low-dimensional population activity in both M2 and A1 is predictive of upcoming and ongoing behavior and is tightly coupled in time.

Given the correlated activity between the principal component projections in A1 and M2, we next identified and quantified shared dimensions of population activity across both areas, using canonical correlation analysis (CCA). Doing so revealed a prominent overlap in latent neural dynamics, with the top canonical dimensions aligning to the motor planning axis observed in both areas independently, while preserving the prominent rotational components associated with motor execution (Fig. 4i; Extended Data Fig. 13a,b). To quantify how each region’s latent dynamics were restructured within the joint manifold, we projected region-specific components (PCs 1-10) into the shared CCA space. In both A1 and M2, dynamics related to motor planning (originally PC3 in each region) were distributed across the top two canonical axes (Extended Data Fig. 13c). To verify that the A1 and M2 geometry was aligned directly via each region-specific latent space and not solely through the top CCA dimensions, we applied Procrustes analysis^54^ to identify the optimal translation, rescaling, and generalized rotation of the low-dimensional population geometries. This analysis revealed that the distance score between the representations was low despite restricting the alignment to only the top dimensions (0.18 +/-0.001 distance score) (Fig. 4j; Extended Data Fig. 14a).

Although strong and temporally correlated alignment between A1 and M2 is consistent with a continuous coupling model, it could instead reflect a common input to both areas rather than continuous communication. To test this possibility, we bilaterally silenced auditory cortex during behavior while recording from M2 (Fig. 4k). During control blocks, M2 activity occupied a stable low-dimensional subspace whose geometry remained strongly aligned with the original intact auditory cortex manifold (Fig. 4l,n). In contrast, silencing auditory cortex induced pronounced deformations in M2’s low-dimensional latent dynamics, affecting dimensions related to both motor planning and motor execution (Fig. 4m,n; Extended Data Fig. 14b). The effect of auditory cortex silencing was not constrained to moments of acoustic feedback, but began immediately after laser activation at trial initiation, consistent with continuous coupling of A1 and M2 (Extended Data Fig. 15a-c). Laser stimulation of auditory cortex in mice not expressing opsins resulted in no perturbation of the low-dimensional M2 manifold, indicating that effects on M2 dynamics cannot be accounted for by behavioral startle (Extended Data Fig. 15d-g). Nor could these effects be explained by overt behavioral differences across conditions, since analyses involved lever presses with equivalent amplitudes (Extended Data Fig. 15h). Finally, changes in M2 activity were not due to stray off-target light, since the behavioral impact of A1 silencing was inconsistent with behavioral changes induced by M2 inactivation (Extended Data Fig. 15i). Together, these data support the idea that continuous input from A1 contributes to sustaining M2’s intrinsic manifold geometry during skilled, acoustic behavior.

Previous work suggests that manifold geometry and local dynamics are heavily shaped by specific behavioral and computational demands^50,55^. If so, we reasoned that the A1 manifold should encode precise motor planning signals in mice trained on a skilled sound-guided acoustic task, but not in mice performing a similar, non-skilled behavior. To test this, we analyzed neural activity in A1 and M2 from mice trained on a non-skilled lever task, in which lever presses produced acoustic feedback and resulted in water reward, but did not require precise motor control to adjust the lever position associated with sound playback or reward (Fig. 4o)^12,30^. These animals exhibited a comparable range of lever-press amplitudes and experienced a similar number of self-generated sounds, allowing for direct comparison of population dynamics across cohorts. In M2, latent dynamics during the non-skilled task remained aligned with low-dimensional dynamics observed during skilled performance (Fig. 4p,r; Extended Data Fig. 16a,b,e,f), with rotational components for motor execution and a separate motor-planning dimension strongly represented across the full-dimensional neural space as revealed by partial least-squares regression (see Methods) (Extended Data Fig. 16h-k). In contrast, A1 of non-skilled mice lacked aligned low-dimensional geometry with the skilled A1 manifold (Fig. 4q,r; Extended Data Fig. 16c,d,e,g). Importantly, motor planning-related dimensions were not detectable in the full-dimensional neural space of A1 in non-skilled mice, while motor execution-related activity persisted (Extended Data Fig. 16h-k). These results indicate that rich signals in motor planning-related neural dimensions emerge only in A1 of mice trained on a skilled acoustic behavior, supporting a model in which cortical manifolds are flexibly updated by behavioral demands to potentially support cross-modal computations.

Together, these results demonstrate that aligned dynamics between A1 and M2 emerge with skilled auditory-motor experience; that latent dimensions are correlated between A1 and M2 throughout behavior; and that ongoing auditory cortex activity is necessary for maintaining the structure of task-related neural dynamics in M2.

## Discussion

Skilled behaviors such as speech and music rely on the brain’s ability to detect performance errors and translate them into precise adjustments in motor output. Yet, the circuit-level mechanisms by which sensory errors are transformed into adaptive changes in behavior remain poorly understood. Here, we establish a mouse model of skilled acoustic behavior and provide evidence that auditory and motor cortices jointly implement this transformation. By combining this novel behavioral paradigm with large-scale electrophysiology and causal perturbations, we identify neural populations that encode sound-guided motor intentions, respond to performance errors, and link error signals to corrective motor plans through shared population dynamics.

Our behavioral paradigm captures key features of real-world sound-guided learning: mice perform goal-directed movements that generate self-produced acoustic feedback, use that feedback to detect performance errors, and adapt their actions rapidly. This provides a novel mammalian model of sound-guided motor learning, which may aid in designing future studies to probe mechanisms underlying flexible motor skill acquisition and performance ^1^. Notably, humans co-opt motor repertoires and auditory perceptual systems to perform non-ethological behaviors such as playing musical instruments. Similarly, despite the artificial nature of the task described here, mice are capable of reusing pre-existing motor skills to acquire the behavior, consistent with the notion that motor systems can flexibly adapt to meet the demands of novel sensorimotor associations.

Behavioral performance was dependent on auditory input, as both tone omission and silencing of auditory cortex disrupted task execution. Brief silencing after movement initiation produced a marked reduction in performance, causing behavior to drop near chance levels, thus underscoring the sensitivity of learned sensorimotor behavior to transient disruption of auditory-motor circuit dynamics. Neurons in auditory cortex responded strongly to performance errors, with distinct populations responsive to overshoot- or undershoot-related feedback. These responses were not purely driven by acoustics: activity during error trials could forecast whether animals adjusted behavior on subsequent trials, consistent with a teaching signal that can be used for corrective motor intentions ^15,56^. A remaining unanswered question is why responses to acoustic error feedback are variable across trials. Our findings argue against these trial-by-trial variations arising due to arousal or attention ^57,58^. Imperfect correction of behavior following acoustic errors has been observed in birds and humans, and may reflect an adaptive neural mechanism for controlling the rate of behavioral adaptation ^26,27^.

Secondary motor cortex (M2) was also needed for successful behavioral adaptation, and error-aligned responses of M2 neurons persistently reflected the type of error that had just occurred (i.e. overshoot or undershoot). M2 dynamics were constrained along a low-dimensional geometry, with errors displacing activity along a dominant planning dimension of activity within this subspace ^59^. These deflections occurred orthogonal to execution-related dimensions, consistent with a selective influence of error signals on future planning rather than ongoing movements. These data support a model of low-dimensional attractor dynamics in motor cortex that can be influenced by sensory feedback during behavior ^28,60^. Our findings are also consistent with recent work showing that orthogonal subspaces in motor dynamics can facilitate computations related to both feedback and control in the same neural populations ^50,61^. While our experiments reveal two brain areas that are necessary for skilled acoustic behavior (A1 and M2) and show activity patterns that correlate with behavioral adaptations, the current findings cannot address whether error-related activity in M2 arrives directly from A1 or through indirect pathways, such as the thalamus or striatum ^36,62^.

A1 activity contained rich information related to motor planning and behavior, including movement amplitude and kinematics. Population activity in auditory cortex was low-dimensional and displayed a geometry in which movement-specific rotations were offset along a third dimension encoding the planned amplitude of upcoming presses, a structure more commonly associated with motor areas, and markedly similar to the manifold geometry that we observed in M2. These findings suggest that auditory cortex may be involved in computations traditionally attributed to motor control, potentially supporting the integration of sensory feedback with predictive motor signals within a predictive processing framework ^16,41^.

Cross-area population analyses revealed a shared low-dimensional latent structure spanning A1 and M2, within which joint neural trajectories reflected motor planning and execution. Silencing auditory cortex disrupted M2 dynamics, suggesting that ongoing input from auditory cortex is required to coordinate adaptive behavior. An interesting question raised by these findings is the extent to which the proposed A1-M2 coupling is robust to partial or transient disruptions. Unilateral or brief perturbations during specific behavioral epochs could help distinguish whether the shared dynamics depend on moment-to-moment interactions or are stabilized by temporal buffering within the underlying circuits. Notably, the shared manifold geometry observed in skilled mice was absent in naïve animals, suggesting that cross-area coordination emerges with learning. These shared population dynamics during skilled behavior are consistent with auditory-motor mirroring during behaviors including vocalizing and music ^8,63^. This framework further positions the planning dimension (i.e. putative output-null space) as a potential substrate for integrating context, error signals, and sensory feedback to guide adaptive motor behavior in real-time ^41,64^.

Cross-area manifold alignment was inferred from non-simultaneous recordings, and future experiments with simultaneous single-trial measurements will be critical to further define the temporal dynamics of this coupling. Future simultaneous A1 and M2 recordings could help determine whether the shared latent geometry reflects trial-by-trial co-variation between areas and reveal the extent of real-time, within-trial coupling that cannot be resolved from non-simultaneous data. A further intriguing direction for future work will be to examine how planning- and movement-related dimensions evolve during within-trial movement corrections, as studies in primate motor cortex suggest dynamic interactions between preparatory and movement-related activity that may reflect continuous updating of motor plans in real time ^65,66^, a question that will require simultaneous single-trial analyses to resolve.

In summary, our findings outline a circuit-level mechanism by which sensory errors are rapidly transformed into updated motor intentions. This framework provides a basis for understanding how sensory feedback may shape ongoing behavior, offering insights into more general principles for flexible, goal-directed control during skilled tasks.

## Acknowledgements

We thank members of the Schneider lab for their thoughtful and valuable comments on this manuscript. We thank Dr. Michael A. Long and Dr. Nicholas J. Audette for their feedback on an early draft of this manuscript, as well as Dr. J Anthony Movshon for general feedback on the project. We thank Jessica A. Guevara and Deanna Garcia for their expert animal care and technical support. This research was supported by the National Institutes of Health (R01DC018802 to D.M.S.; F31DC021868 & T32NS086750 to G.W.Z.); a Career Award at the Scientific Interface from the Burroughs Wellcome Fund (D.M.S.); fellowships from the Searle Scholars Program, the Alfred P. Sloan Foundation, and the McKnight Foundation (D.M.S. and A.H.W.); and an investigator award from the New York Stem Cell Foundation (D.M.S.). D.M.S. is a New York Stem Cell Foundation - Robertson Neuroscience Investigator.

## Author contributions

Conceptualization, G.W.Z. and D.M.S.; investigation, G.W.Z.; data collection, G.W.Z., W.Z., B.E.H.; data analysis, G.W.Z.; writing – original draft, G.W.Z. and D.M.S.; writing – review & editing, G.W.Z., W.Z., A.H.W., D.M.S.; supervision, A.H.W., D.M.S.; funding acquisition, D.M.S.

## Methods

### Animals

All experimental protocols were approved by New York University’s Animal Use and Welfare Committee. Male and female wild-type (C57BL/6) or VGAT-ChR2-EYFP mice were purchased from Jackson Laboratories and subsequently housed and bred in an onsite vivarium. Mice were co-housed with littermates during training and singly housed following craniotomy surgery. Water restricted mice were given ~1 mL of water per day via task-based rewards and supplements to maintain 80–90% of the animal’s pre-restriction body weight. Animals of both sexes were used for experiments (12 females, 9 males). All experimental mice were 3–4 months old and were kept on a reverse day/night cycle (12h day, 12h night).

### Craniotomy and head-post implantation surgeries

For all surgical procedures, mice were anaesthetized under isoflurane (3% induction; 1.5%–2% maintenance in O2) and placed in a stereotaxic holder with non rupture, zygoma ear cups (Kopf Instruments, models 963 and 1721) with a heating pad to maintain and monitor body temperature (Harvard Apparatus). The scalp was disinfected with 70% ethanol and betadine. Mice were then injected with 150 µl of local anesthetic under the scalp (0.25% bupivacaine hydrochloride saline solution, Sigma-Aldrich B5274-1G). Eyes were covered with Vaseline. Skin was then removed over the top of the head, the surrounding edge of the exposed cranium was covered with tissue adhesive (Vetbond, 3M), and a custom laser cut Y-shaped titanium headpost (1.5 mm thickness, 120° angle, 3 branches, sendcutsend.com) was attached to the skull using dental cement (Lang Dental). Following surgery mice were treated with an analgesic (Meloxicam, SR), placed on a heating pad, and allowed to recover for 3-5 days prior to training. Following training and prior to electrophysiology, a small craniotomy (~1.5mm diameter) was made using a dental drill (Foredom, model 1474) and micro drill bit (0.9 mm head diameter, Fine Science Tools, No. 19007-09) to expose either the auditory or secondary motor cortex. A second craniotomy (~0.5mm diameter) was made away from the recording site over the parietal bones for implantation of a grounding bone screw (Fine Science Tools, No. 19010-11). Exposed craniotomies were covered with a silicone elastomer (World Precision Instruments, Kwik-Sil) and the mouse was allowed to recover in its home cage prior to electrophysiology.

### Behavioral task and data collection

A custom-designed lever (7 cm long, 3D-printed using Formlabs Form2) was mounted to the post of a rotary encoder (US Digital) 5 cm from the lever handle. A magnet (CMS magnetics) was mounted to the bottom of the lever, which was positioned 5 cm above a larger static magnet which established the lever resting position and provided light and adjustable movement resistance. The lever handle (top) was positioned adjacent to the front of a tube (Custom, 3D-printed using Formlabs Form2), which held mice directly below two plate clamps to secure the mouse headpost. Lever and mouse apparatus were constructed with Thorlabs components. A water spout, controlled by a solenoid valve (The Lee Company), was positioned in front of the mouse to allow for reward delivery. Licking was measured using a custom-mounted (3D printed using Formlabs Form2) IR-beam emitter and receiver. The IR signal was pre-processed using a custom printed circuit board to generate a binary TTL signal.

Behavioral data signals for licking and lever movement were acquired by a single Arduino UNO running a custom script written in Arduino IDE (sampled >3kHz) that communicated in real time with custom written MATLAB software (Mathworks, PsychToolBox) via serial communication to control the virtual environment of the experiment. An additional Arduino UNO was used to deliver tones and TTL pulses with sub-millisecond delays. We recorded sounds during some experiments using an ultrasonic microphone (Avisoft, Model # CM16/CMPA-P48) positioned 1 cm from the lever to confirm that the lever produced negligible noise (<1 dB SPL) and that experimenter-controlled sounds were delivered at a consistent volume (70dB SPL). All training was performed in a sound-attenuating booth (Gretch-Ken) monitored via three IR video cameras (AAK CA20 600TVL 2.8MM).

During behavior, mice were head-fixed to the behavioral apparatus and presented with the lever and lick-spout after quiet acclimation. Mice were then allowed to engage with the behavioral task at will. Lever movements into the target zone (see results) elicited a small water reward (5-10uL) when the lever returned to home position. Auditory feedback in the form of a 16kHz tone (50ms duration, 70 dB) was delivered on all trials when the lever entered the zone, and in the form of an additional 8kHz tone (50ms duration, 70 dB, start time delayed by 50 ms relative to zone exit point) if the lever exceed the bounds of the zone. Entry and exit tones produced subtle broadband clicks at the onset and offset, which we found was important for achieving expert behavior. Mice were initially trained (~7-10 days) with a single 16kHz frequency elicited at the apex of any lever movement that entered a large target zone (~4x the size of the final expert zone size). Once proficiency with the apparatus was achieved, mice were introduced to the full task structure with three smaller zones (see results). The exact placement of the zones was slightly different (+/-2mm) for each mouse depending on their body size and average pressing range early in training. Mice received between 15 and 25 sessions of training over 15-20 days before electrophysiology and/or optogenetic stimulation experiments, with either one or two sessions per day. The datasets for non-skilled behavior have been described previously ^12,30^. Here, we applied new analytical approaches to these neural recordings.

### Electrophysiology recording and spike curation analysis

Following behavioral training, we used phase maps generated via intrinsic optical signal imaging (IOS) to guide surgical opening of a craniotomy above the center of primary auditory cortex. To target the secondary motor cortex (M2), stereotaxic coordinates (1.0-1.5mm AP, 0.5-0.7 mm ML from bregma) were used. During electrophysiological recordings, mice were positioned in the behavioral apparatus and a 128-channel electrode (128AxN, Masmanidis Lab) was lowered into either the auditory or secondary motor cortex orthogonal to the pial surface (Yang et al., 2020). The electrode was connected to a digitizing head stage (Intan Technologies) and electrode signals were acquired at 30 kHz, monitored in real time, and stored for offline analysis (Open Ephys Acquisition Board and GUI). The probe was allowed to settle for at least 20 minutes prior to behavior. After behavior, mice were permitted to rest without access to the lever for 10 mins before a set of tones (including stimuli identical to those used during the task) were played. The total length of electrophysiological recordings lasted approximately 45-60 minutes.

After recording, electrical signals were processed and the action-potentials of individual neurons were sorted using Kilosort4^67^ and manually curated in Phy2.0. During manual curation, clusters were evaluated based on estimated refractory-period contamination, waveform principal component structure, interspike interval histograms, and stability of firing rates throughout the behavioral session. Units exhibiting substantial multi-unit contamination, unstable recordings, or nonphysiological waveforms were excluded from further analysis. Across 3,022 curated units, the estimated contamination was 5.8% (median; IQR, 0.8-15.8%). All subsequent analyses were performed using the manually curated spike times assigned to each unit.

### Intrinsic optical signal imaging

Neural recordings and optogenetic silencing experiments were targeted to the auditory cortex using transcranial intrinsic optical signal imaging (IOS). During the procedure, the region of the skull roughly above the auditory cortex was slightly thinned using a dental drill, and cleared for optical access by applying cyanoacrylate. Imaging was performed under isoflurane anesthesia (3% induction; 1-1.5% maintenance in O2). The brain surface was illuminated with red light using a 680-nm Fiber-Coupled LED at ~2mW (Thorlabs M680F4). Images were acquired through a 4X air objective (NA 0.10, Olympus) using a CMOS camera (PCO Edge 3.1; 16 bit; 2048×1536 pixels, binned down to 512×384 pixels; 50 Hz frame rate, binned down to 10Hz). The imaging plane was set 400-600 μm below the cortical surface. A still image of the cortical surface was acquired using green light from a white LED (Thorlabs MCWHL5) bandpass filtered (469/35nm) to visualize the blood vessel pattern.

The primary auditory cortex was identified through tonotopic mapping using a temporally periodic acoustic stimulation consisting of sequences of single pure tones of ascending or descending frequencies (20 tones per sequence, 2 to 40kHz exponentially separated, 75dB SPL, 50ms tone duration, 500ms onset-to-onset interval). Tone sequences were delivered from a free-field speaker 10cm above the mouse and repeated 40-60 times (0.1Hz). To compute tonotopic maps, the time course of each pixel was first high-pass filtered using a moving average (10 second window). Next, a Fourier transform was computed to extract the phase and the power of the frequency component at the frequency of acoustic stimulation (0.1 Hz). The phase indicates the sound frequency driving the response of a pixel, and the power indicates the strength of its response. To account for the hemodynamic delay and compute maps of absolute tonotopy, the response time to ascending sequence of tones was subtracted from the response time to the descending sequence. From these maps of absolute tonotopy, equally spaced iso-frequency contour lines were extracted, color-coded for sound frequency, and overlaid on top of the image of the blood vessel pattern (Extended Data Fig. 5a). The primary auditory cortex was identified based on the established tonotopic representation of mouse auditory cortex, by locating the axis between the most posterior low-frequency peak and the large, central high-frequency peak ^68^. Image acquisition and analysis software were custom written in MATLAB.

### Optogenetic silencing experiments

Real time optogenetic stimulation of either bilateral auditory or secondary motor cortex was accomplished via TTL control (custom MATLAB and Arduino scripts) of an all solid-state 473nm blue laser (MBL-III-473/1~100mW, Opto Engine LLC). Bifurcated fiber cables (ThorLabs Ø200 µm Core, 0.39 NA, SMA905 to Ferrules) were used for light delivery. The laser had a diameter of 1 mm at the surface of the cranium, centered and guided by either phase maps generated via IOS (primary auditory cortex) or stereotaxic coordinates (secondary motor cortex). During optogenetic perturbation trials, mouse initiated presses resulted in closed-loop activation of the laser (~15-20mW) for the duration of a press (mean duration 406ms +/-0.02). Additionally, the laser was deactivated if a mouse press lasted longer than 3 seconds (less than 1% of trials).

### Decoding analyses

Population decoding analyses were performed on spike count data using time bins as specified in each figure legend. To ensure balanced data across conditions, trials were randomly subsampled to the minimum number of trials available per condition within each animal. Data were normalized by z-scoring each neuron’s activity across all conditions. Decoding was implemented using a multiclass linear support vector machine (SVM) classifier employing error-correcting output codes (*fitcecoc*, MATLAB Statistics and Machine Learning Toolbox). For each decoding iteration, 80% of the data were randomly assigned for training and 20% for testing. This process was repeated 1000 times with random subsampling to account for trial variability and to estimate decoding reliability. To assess chance performance, trial labels were shuffled and decoding was repeated with identical parameters. Decoding accuracy was computed as the fraction of correctly classified trials in the test set and averaged across iterations for each animal. All analyses were implemented in custom written MATLAB scripts.

### Population dynamics analyses

#### Principal Component Analysis (PCA)

Neural activity from auditory cortex (A1) and secondary motor cortex (M2) recordings (non-simultaneous) was characterized by performing principal components analysis (PCA) on trial averaged peri-stimulus time histograms (PSTHs) pooled across animals. Only neurons with a minimum of 10 trials per condition were included. PSTHs were binned at 10 ms resolution and smoothed with a Gaussian kernel (σ = 20 ms) for all population dynamics analyses. Each neuron’s trial-averaged PSTHs across all conditions were concatenated to form a neurons x CT_c_ matrix, where C is equal to the number of conditions and T_c_ the number of time points per condition. Prior to dimensionality reduction, neural responses were z-scored per neuron by subtracting the mean and dividing by the standard deviation across all time points and conditions.

Low-dimensional representations of the population data were obtained using PCA. To estimate the variability and confidence bounds of the low-dimensional projections across trials, we implemented a resampling procedure consisting of 1000 iterations of 50-50 random splits of trials within each condition, generating matched training and testing datasets. For each run, the principal component projection matrix was calculated on one partition of the data and then applied on the second partition to yield a low-dimensional embedding in the latent space. The mean across resampling runs was taken as our estimate of the low-dimensional trajectory and the central 95 percentiles across runs was used to approximate the confidence intervals (CIs).

For analyses involving error projection onto the M2 ‘planning-axis’ (Fig. 3) and to assess variability in error-related deflections across the neural population, a bootstrap analysis was applied (1000 iterations). During each run, PCA was performed on a resampled population of neural PSTHs computed on pre-trial neural activity (−540 ms to −290 ms preceding trial onset) for all trials of a given amplitude condition to generate ‘planning-axis’ neural loadings. Trial-averaged error PSTHs (−125 ms to +125 ms surrounding time of acoustic error) were then projected onto the top principal component and neural loadings derived from the pre-trial activity. Each error PSTH was normalized using the training data mean and standard deviation before projection onto the pre-trial PC1. Error bars represent approximate confidence intervals given by the central 95 percentiles resulting from resampling of the neural population. For this analysis, undershoot aligned PSTHs were uniformly advanced by +75 ms relative to original apex alignment to account for the increase in activity before time zero shown in Figure 2l (see statistical methods below).

#### Canonical Correlation Analysis of shared population dynamics across brain regions

To quantify the shared latent dynamics between primary auditory cortex (A1) and secondary motor cortex (M2), we performed canonical correlation analysis (CCA). To reduce dimensionality and denoise the data prior to CCA, we used PCA projections of microzone aligned neural activity derived from a pseudopopulation across animals (as above, but without resampling). To ensure stable PCA axes for CCA, dimensionality reduction was performed on trial averaged PSTHs computed using the full set of trials for each microzone. In this analysis, our aim was to characterize population-level structure rather than evaluate generalization performance across trials. Notably, PCA on microzone neural activity per the subsampling approach above yielded qualitatively similar results to the non-subsampled version in the top principal components, supporting the stability of the dominant axes of variance used in subsequent analyses.

To select the optimal number of principal components (PCs) for each region, we ran a repeated analysis over PC dimensionalities (5 to 20 PCs for both A1 and M2). For each PC pair, we performed leave-one-out cross-validation. On each fold, PCA and CCA were fit on all but one condition (*canoncorr*, MATLAB), and the held-out condition was projected into the learned canonical subspace. The correlation coefficient (i.e., the correlation along the dominant shared dimension) was then computed on the held-out condition across regions (*corr*, MATLAB). This process was repeated for all microzones, and the mean cross-validated correlation was stored. The PC dimensionality pair yielding the highest mean canonical correlation was selected for all further analyses (A1: 15 PCs, M2: 17 PCs). Predictive generalization between the full dimensions of the shared subspace between A1 and M2 was evaluated independently using further cross-validated regression analyses (see below). Importantly, results were qualitatively robust across a broad range of PC dimensionalities, indicating that the specific number of PCs selected for each region did not substantially impact the structure or strength of A1-M2 alignment. For this reason, CCA analysis of skilled versus non-skilled neural manifolds depicted in Extended Data Figure 16 utilized a fixed number of 20 PCs per neural population prior to alignment (Extended Data Fig. 16f,g).

With the optimal number of PCs determined, we performed CCA on the reduced data to identify low-dimensional subspaces in A1 and M2 that maximally covaried across conditions. PCA scores from each region (samples x PCs) were input into MATLAB’s *canoncorr* function, which returned transformation matrices projecting each region’s activity into the shared canonical space. To evaluate the reliability and predictive utility of the shared latent trajectories, we conducted a second round of leave-one-condition-out cross-validation in the top four canonical dimensions. On each fold, CCA was fit on the training conditions, and a linear regression model was trained to predict M2 canonical activity from A1 canonical activity. This model was then applied to the held-out condition, and predictive performance was quantified using the coefficient of determination (R^2^) between predicted and actual M2 activity in the shared space. The mean cross-validated R^2^ across conditions was used to assess the generalizability of the shared subspace between A1 and M2 (mean R^2^ = 0.96). To visualize population dynamics, we projected trial-averaged activity from both A1 and M2 onto the top three canonical dimensions for each condition (Fig. 4). Additionally, we plotted the time courses of individual canonical dimensions to examine temporal alignment of trajectories between regions (Extended Data Fig. 13a,b).

#### Procrustes Alignment of neural population trajectories

To quantify the geometric similarity of low-dimensional neural representations between brain regions and/or recordings, we performed Procrustes alignment on trial-averaged neural activity pooled across animals. This approach captures the similarity in the shape of neural representations while allowing for rotation, scaling, and translation differences between regions.

For each of the press amplitudes, PSTHs were computed across conditions for each neuron by alignment to trial onset (−500 ms to +1000 ms), binned at 10 ms resolution and smoothed with a Gaussian kernel (σ = 20 ms). Only neurons with a minimum of 10 trials per condition were included. The data for each condition and region was organized into matrices of dimensions neurons x CTc (where C is equal to the number of conditions and Tc the number of time points per condition) then reshaped and concatenated across conditions for subsequent analysis. For measurements taken of aligned low-dimensional geometry, dimensionality was fixed at 3 principal components for each population prior to the alignment. Neural responses were z-scored across time and conditions within each neuron and then projected into a low-dimensional latent space using a subsampling procedure described below for 20 data splits. For each press amplitude condition, trials were randomly split 20 times into equal halves (50-50) across regions (i.e., 50% of A1 trials compared to 50% of M2 trials). For each iteration, PCA was performed separately on PSTHs calculated using the subsampled trials per region, and Procrustes alignment was then applied following the procedure described below.

For each region, PCA-projected trajectories from all conditions were concatenated into a single matrix of size timepoints x dimensions. The Procrustes transformation was then computed to optimally align the representations, allowing for rotation, scaling, and translation (procrustes, MATLAB). The reference region used in each specific analysis is noted in the figure legends accompanying the results. The resulting Procrustes residual distance score quantified the degree of alignment as the sum of the squared differences between the shapes, with a lower score indicating greater similarity in the geometry of latent dynamics (0 = perfect fit, 1 = maximum dissimilarity). Aligned trajectories were visualized in 3D space (PCs 1-3), with different colors/shades representing trajectories from aligned A1 and M2 data (Fig. 4).

To assess the statistical significance of the observed alignment score between A1 and M2 dynamics reported in Figure 4j, we generated a null distribution by disrupting the temporal structure of the M2 latent trajectories. Specifically, after PCA projection, we shuffled the time bins of each condition’s M2 trajectory, thereby preserving the overall distribution and dimensionality of activity while eliminating meaningful temporal dynamics. These temporally shuffled trajectories were then concatenated across conditions and aligned to the unshuffled A1 trajectories using the same Procrustes procedure described above. This process yielded a distribution of alignment scores expected by chance across the 20 data splits. The alignment score obtained from the real data was compared to this null distribution to determine whether the observed similarity between A1 and M2 dynamics exceeded what would be expected from random temporal structure alone (Extended Data Fig. 14a).

To assess the similarity of latent neural representations between A1 and M2 under conditions of auditory cortex (AC) silencing and intact AC (Fig. 4), and to evaluate the stability of these representations across trials, we implemented a subsampling procedure. For each condition, trials were randomly split 20 times into equal halves (50-50) within and across regions (i.e., 50% of A1 trials compared to 50% of M2 trials during AC intact, and 50% of A1 trials compared to 50% of M2 trials during AC silencing). For each iteration, PCA was performed separately on PSTHs calculated using the subsampled trials per condition and region, and Procrustes alignment was then applied following the procedure described above. The resulting distribution of Procrustes residual scores, reflecting alignment quality across data splits, is reported in Figure 4n. To account for the reduction in trial count during optogenetic silencing blocks, all analyses in this case were conducted using neural activity during presses to each of the 3 original zones (rather than the 9 microzones). A similar procedure to the above was performed to assess the similarity of geometry across non-skilled and skilled datasets reported in Figure 4p-r.

#### Partial least-squares regression

To quantify the detectability of skilled-M2 planning-related neural activity across all other neural recordings, we first concatenated pre-trial aligned PSTHs (−1000 ms to −300 ms) across the 3 zones and z-scored each neuron’s activity across conditions. PCA was applied to the concatenated PSTHs, and the first principal component (PC1) was defined as the “planning-axis”. M2 neural activity was then projected onto this axis to generate an M2 reference planning projection in PC space.

To assess the robustness of this signal across recordings, we performed partial least-squares (PLS) regression (*plsregress*, MATLAB). For each of 20 data splits, a subset of trials from all conditions was used to calculate neural PSTHs aligned to trial start (−500 ms to +1000 ms) and train a PLS model predicting the original pre-trial M2 reference planning projection from population activity directly in the full dimensional neural space (i.e. without PCA pre-processing). The goal was therefore to quantify the presence of the skilled-M2 defined planning-axis across all other neural recordings (non-skilled M2, non-skilled A1 etc.). Following training, the model was then tested on the other 50% of held-out trials. Two metrics quantified performance: the Pearson correlation between predicted and reference projections, and the fraction of variance explained (Extended Data Fig. 16h-k). Both metrics reflect the strength and presence of planning-related population dynamics across neurons and conditions. An equivalent analysis was performed using neural PSTHs initially calculated on activity −500 ms to +1000 ms surrounding trial initiation to define an “execution-axis” (PC1) from skilled-M2 neural activity and to quantify the detectability of M2 execution-related dimensions across all other recordings using the same PLS procedure described above (Extended Data Fig. 16h-k).

### Trial-duration normalization

For the analyses shown in Extended Data Figures 8 and 10, peri-stimulus time histograms (PSTHs) were computed using a common time-binning procedure to account for small differences in trial duration across the three conditions. A common analysis window was defined across all 3 trial types using the 90th percentile of trial durations, extending from −200 ms before trial onset to the percentile-based trial end. This window was discretized into a fixed number of uniformly spaced bins (200 total bins), and the resulting global bin edges were applied identically across all trials. For each neuron and trial, spikes within the analysis window were binned and converted to firing rates by dividing bin counts by the bin width. Trial-specific firing rates were then averaged across trials to obtain the mean PSTH for each neuron. Mean PSTHs were then smoothed with a Gaussian kernel of width 30 bins.

### Statistical analyses

All sizes of animal cohorts or individual experiments are denoted by a capital N, while cell population sizes are denoted by a lowercase n. All error bars represent either standard error (standard deviation divided by square root of the size of the analysis population) or approximate 95% confidence intervals and are specified on a panel-by-panel basis. A two-sided, non-parametric Wilcoxon rank-sum test (*ranksum*, MATLAB), was used for most statistical comparisons reported throughout including comparisons between differing subpopulations, decoding results for real and shuffled data, and Procrustes alignment scores across regions/conditions. To identify neurons with significant increases in responsiveness to stimuli relative to baseline (i.e., entry tone, exit tone, etc.), an equivalent one-sided test was used. For comparisons of the same mice across behavioral conditions (Fig. 1) and behavioral perturbation data (Fig. 2, Fig. 3), a parametric paired t-test (two-sided) was used (ttest, MATLAB). Statistical significance on figures was denoted as *p < 0.05, **p < 0.005, ***p < .001. Parentheticals with descriptive statistics throughout the text and figure legends represent mean values +/-standard deviation.

For data reported in Figure 2, given that the precise timing of omission encoding varied from one trial to the next due to lever kinematics (and thus is not perfectly consistent for non-simultaneous neurons across recordings), we applied a sliding window analysis on neural spike counts (50 ms bins, advanced in 10 ms steps) spanning −500 ms to +500 ms around the undershoot apex to identify neurons with significant increases in omission-related activity. This analysis revealed a temporal concentration of omission-related responses as early as 75 ms before our artificially labeled ‘undershoot error’ time, and shortly following the expected tone onset on correct trials (Figure 2l). We therefore advanced all M2 population projections from undershoot trials shown in Figure 3 by +75ms during analysis to better estimate the time of error signaling (Fig. 3o,p).

For data reported in Figure 2h,p, a two-sided permutation analysis was performed to generate a null distribution of modulation indices favoring each corrected/non-corrected condition to perform a two-sided statistical test. First, data for all neurons recorded was organized into two separate neurons-by-trials matrices consisting of individual spike counts calculated in equal sized time windows for each trial type. Then, for each individual neuron a modulation index (MI) was calculated according to the average spike counts (C) in each condition during the window of interest (C_1_-C_2_)/(C_1_+C_2_). For 1000 permutations, the spike counts all neurons from each condition were randomly shuffled and the MI was recalculated to generate the null distribution; approximate 95% confidence intervals were estimated as 2.5-97.5th percentiles of all permutations. Therefore, a neuron was considered to statistically favor a condition if its real index was found to be in either the 2.5 or 97.5 percentile of the null distribution.

---

**Extended Data Fig. 1.**
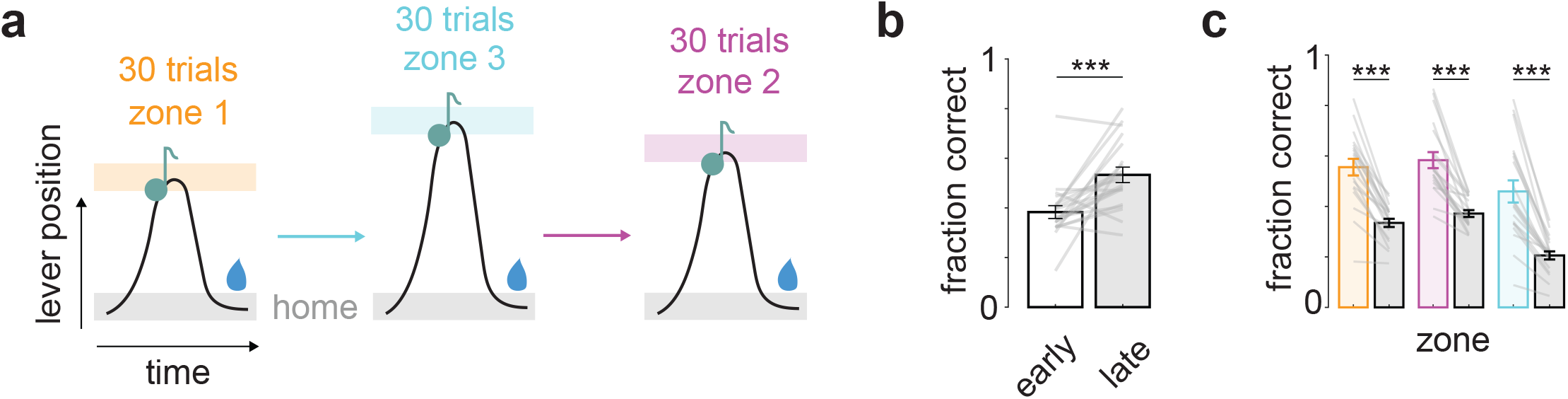
Mice perform presses to 3 different zone locations. **a**, Schematic depicting zone location shift every 30 trials. Zone size and acoustic cues remain constant. **b**, Fraction of trials that are correct across early versus late training sessions, (N=20, paired t-test, ***p<0.001). **c**, Fraction of correct trials per individual zone during late training sessions compared to chance levels (gray), (N=20, paired t-test, ***p<0.001).

**Extended Data Fig. 2.**
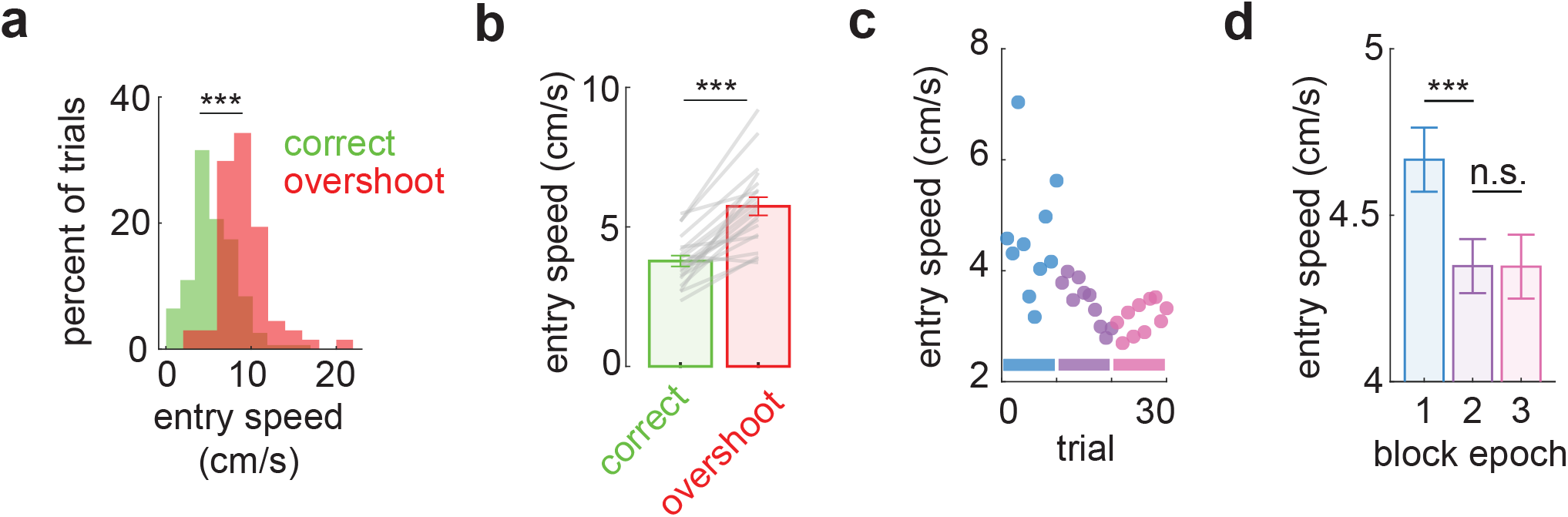
Mice anticipate zone location through experience. **a**, Zone entry speed during all correct and overshoot trials for an example mouse behavioral session, (ranksum, ***p<0.001). **b**, Mean zone entry speed quantified across all mice for correct and overshoot trials (N=20, paired t-test, ***p<0.001). Error bars represent SEM. **c**, Mean zone entry speed across trials in a block for an example mouse, (blue: trials 1-10, magenta: trials 11-20, and pink: trials 21-30). **d**, Mean zone entry speed across block epochs for all mice (N=20, ranksum, ***p<0.001; n.s. not significant).

**Extended Data Fig. 3.**
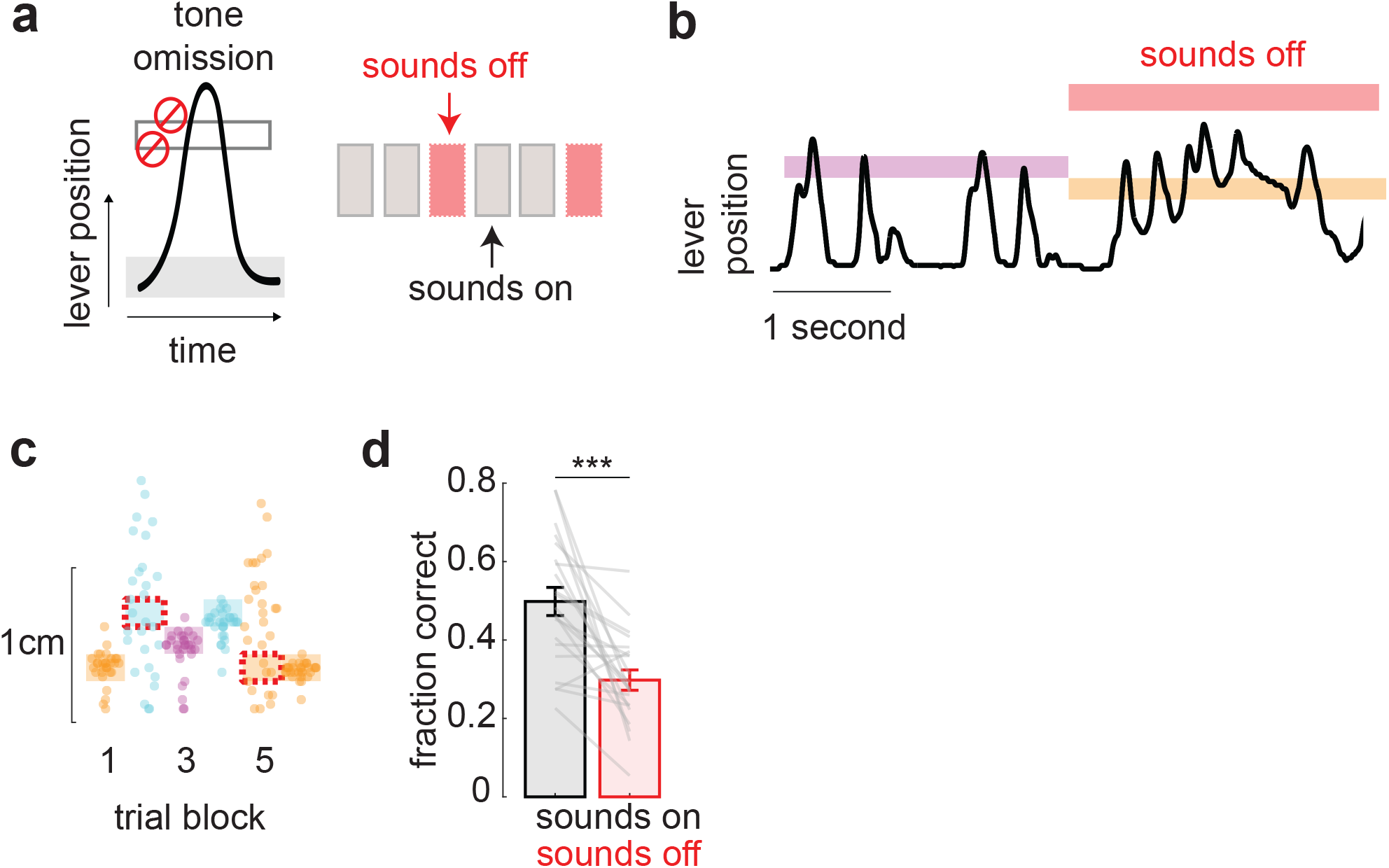
Mice rely on acoustic cues to perform the behavior. **a**, Schematic depicting tone omission experiments. Every 3^rd^ block, both tones bordering the target zone are removed for the duration of the entire block. **b**, Example lever trace during transition period to an omission block. **c**, Example behavior during 6 trial blocks. Individual trial press peaks are plotted as scattered dots. Shaded squares represent the confines of the zone presented for 30 trials at a time. Red hashed squares represent blocks during which acoustic cues are omitted. **d**, Mean performance (fraction correct) following a correct trial (i.e., two correct presses in a row) shown for all mice during control and tone omission blocks (N=21, paired t-test, ***p<0.001).

**Extended Data Fig. 4.**
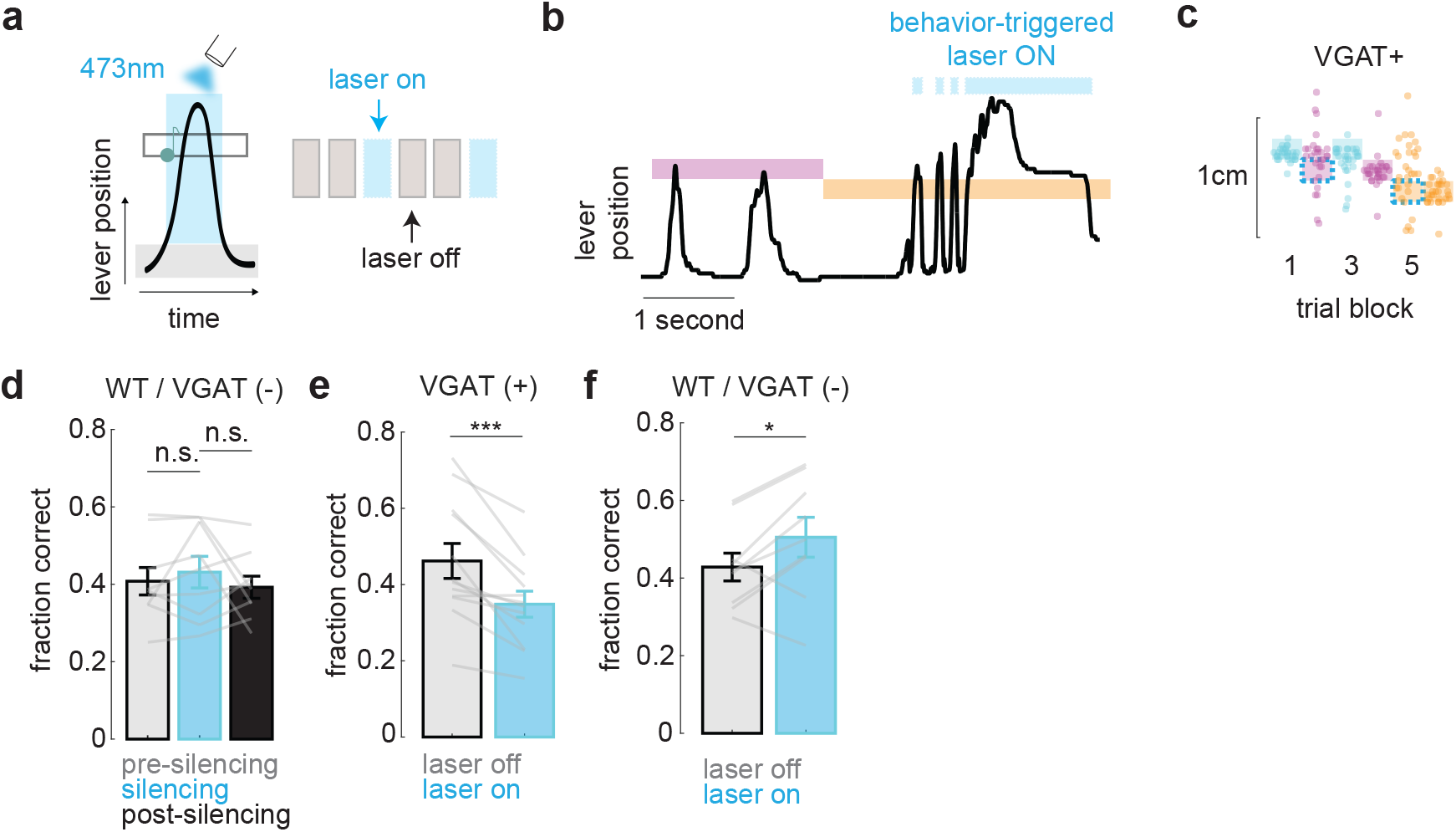
Performance decreases with auditory cortex silenced only in VGAT(+) mice. **a**, Schematic depicting closed loop activation of 473nm laser during self-initiated trials. Every 3^rd^ block, the laser was activated on every trial in that block. **b**, Example lever trace during transition period to an optogenetic perturbation block. **c**, Example behavior during 6 trial blocks. Individual trial press peaks are plotted as scattered dots. Shaded squares represent the confines of the zone presented for 30 trials at a time. Cyan hashed squares represent optogenetic perturbation blocks. **d**, Mean performance (fraction correct) shown for wild-type (WT)/VGAT(−) mice during AC optogenetic perturbation blocks and the control blocks surrounding the perturbation blocks, (N=9, paired t-test; n.s. not-significant). **e**, Mean performance (fraction correct) following a correct trial shown for all VGAT(+) mice during control and AC optogenetic perturbation blocks (N=12, paired t-test, ***p<0.001). **f**, Mean performance (fraction correct) following a correct trial shown for all wild-type (WT)/VGAT(−) mice during control and AC optogenetic perturbation blocks (N=9, paired t-test; *p<0.05).

**Extended Data Fig. 5.**
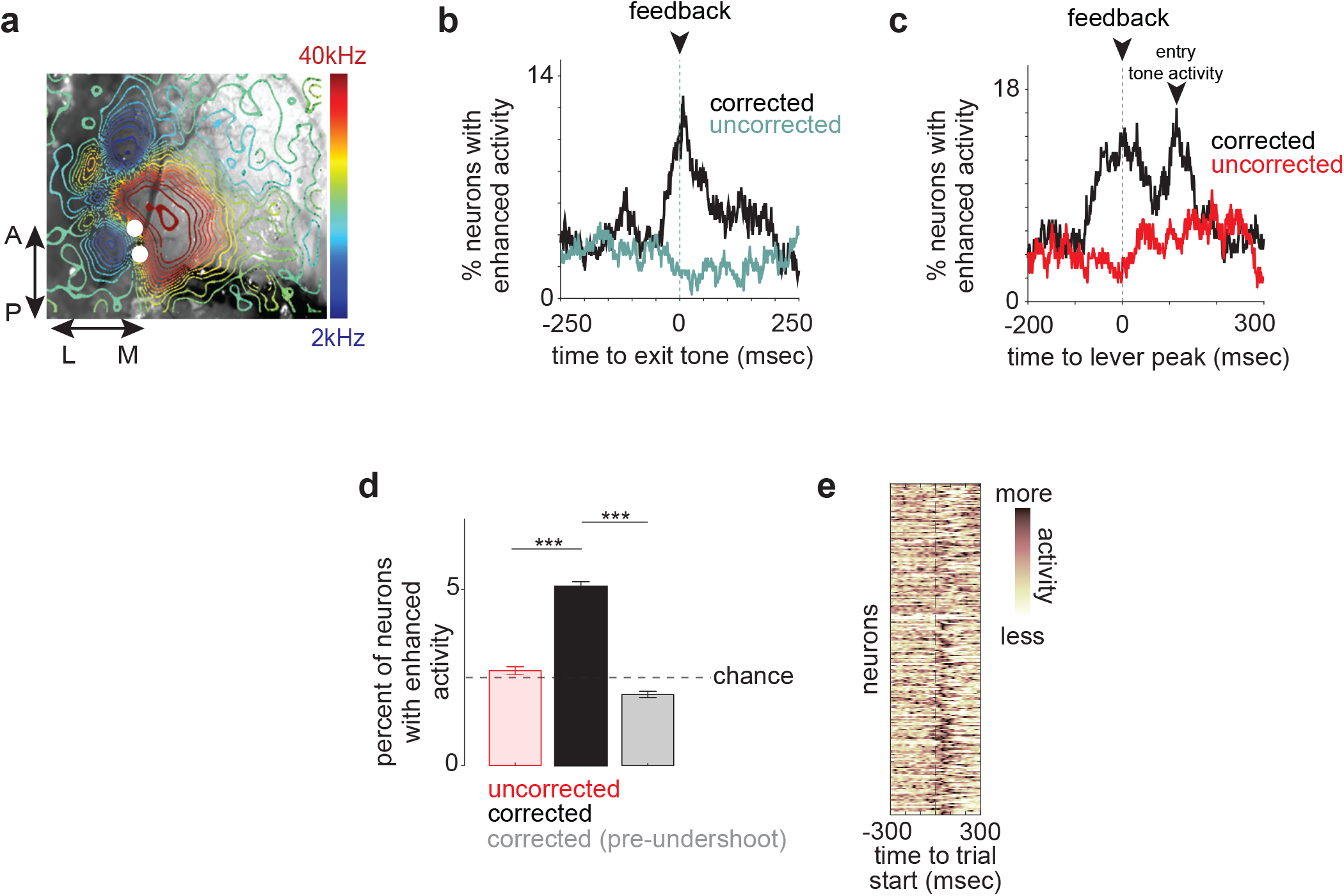
Auditory cortex targeted electrophysiology reveals error encoding populations. **a**, A1 targeting using intrinsic optical signal (IOS) imaging. White dots represent electrode implantation sites during electrophysiology. **b**, Sliding window (50ms window advanced in 1ms steps) analysis in which spike counts are recomputed across each condition and a two-sided permutation test (see Methods) is applied to identify the percent of neurons signaling next-trial corrections during overshoot trials. **c**, Same analysis as in (**b**), but data shown for undershoot trials. **d**, Two-sided permuation test (see Methods) identifying percent of neurons signaling next-trial correction following an uncorrected undershoot using neural activity −75 ms to +75 ms surrounding initial undershoot apex. Control (“pre-undershoot”) indicates equivalent analysis using activity −350ms to −200ms prior to undershoot apex on same trials. Hashed line represents 2.5% chance level (ranksum, ***p<0.001). **e**, Trial averaged PSTHs of omission responsive neurons recorded from A1 depicted in Figure 2n, now aligned to trial initiation of correct trials (i.e. non-undershoot event trials). Each neuron’s PSTH normalized to occupy range of 0-1.

**Extended Data Fig. 6.**
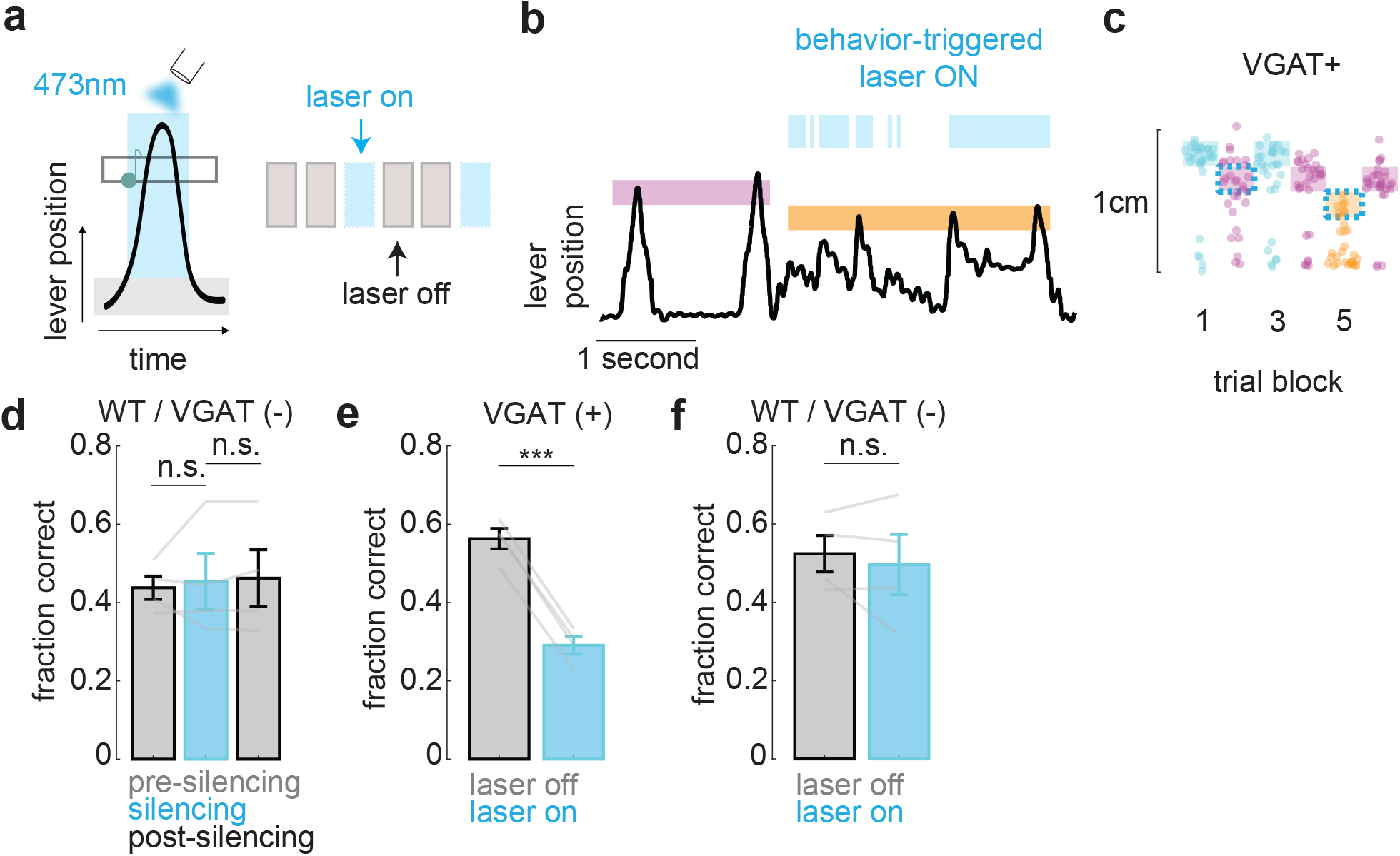
Performance decreases with M2 silenced. **a**, Schematic depicting closed loop activation of 473nm laser during self-initiated trials. Every 3^rd^ block, the laser was activated on every trial in that block. **b**, Example lever trace during transition period to an optogenetic perturbation block. **c**, Example behavior during 6 trial blocks. Individual trial press peaks are plotted as scattered dots. Shaded squares represent the confines of the zone presented for 30 trials at a time. Cyan hashed squares represent optogenetic perturbation blocks. **d**, Mean performance (fraction correct) shown for wild-type (WT)/V-GAT(−) mice during M2 optogenetic perturbation blocks and the control blocks surrounding the perturbation blocks, (N=4, paired t-test; n.s. not-significant). **e**, Mean performance (fraction correct) following a correct trial shown for all VGAT(+) mice during control and M2 optogenetic perturbation blocks (N=4, paired t-test, ***p<0.001). **f**, Mean performance (fraction correct) following a correct trial shown for all wild-type (WT)VGAT(−) mice during control and M2 optogenetic perturbation blocks (N=4, paired t-test; n.s. not-significant).

**Extended Data Fig. 7.**
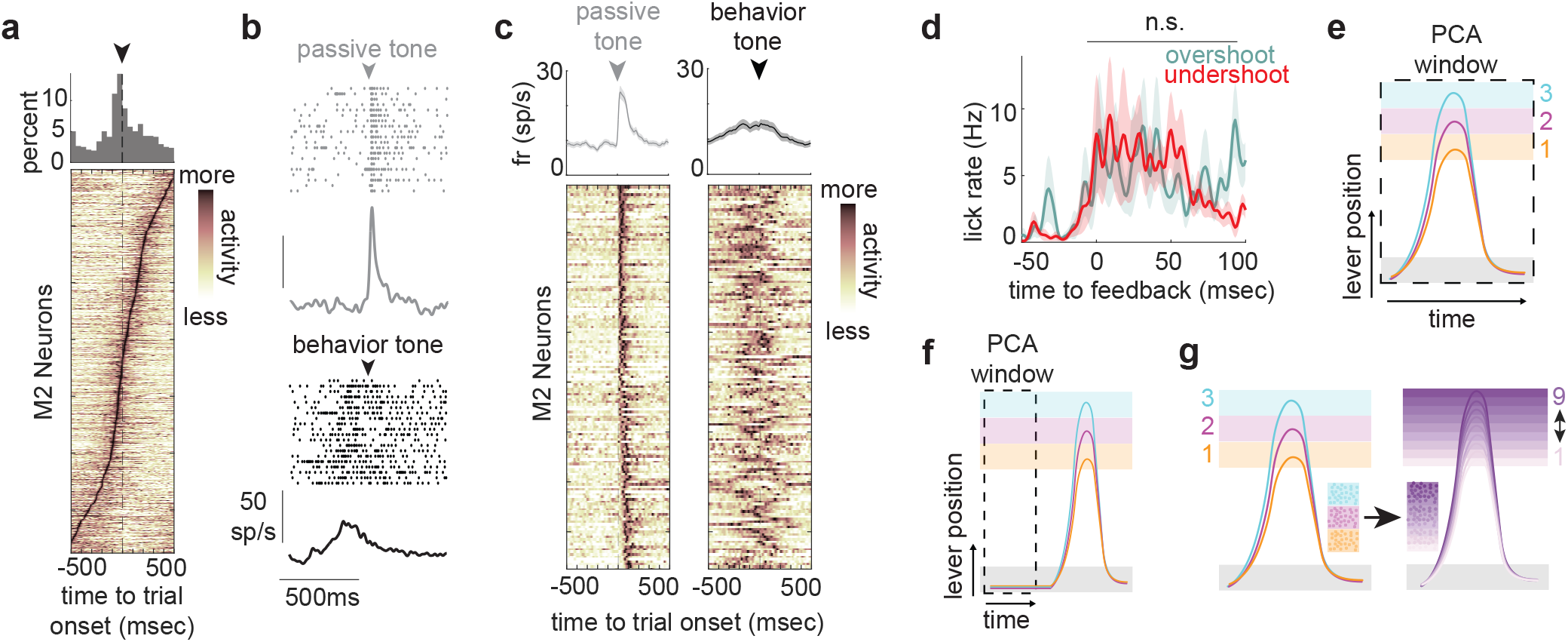
M2 neurons are active during behavior and rapid onset sound responses are gated off. **a**, Bottom: full M2 neural data set (n=676, N=9). Peri-stimulus time histograms (PSTHs) shown as rows in heatmap normalized to occupy range 0-1, computed on neural activity −500ms to 500ms surrounding trial start on correct trials. PSTHs organized by time of peak firing rate across window of analysis. Top: histogram of individual neuron peak times during behavior. **b**, Example M2 neuron response to passive and active presentation of the 16kHz entry tone. **c**, Neural responses to the entry tone heard passively and during behavior for the full population of passively sound responsive M2 neurons. **d**, Average licking rate PSTHs aligned to overshoot and undershoot errors. (ranksum, n.s. not-significant). **e**, Schematic depicting activity window used for PCA training and results depicted in (**Figure 3g**). **f**, Schematic depicting activity window used for PCA training and results depicted in (**Figure 3h**) and (**Figure 3i**). **g**, Schematic depicting target zones partition into 9 equally sized microzones covering the same range.

**Extended Data Fig. 8.**
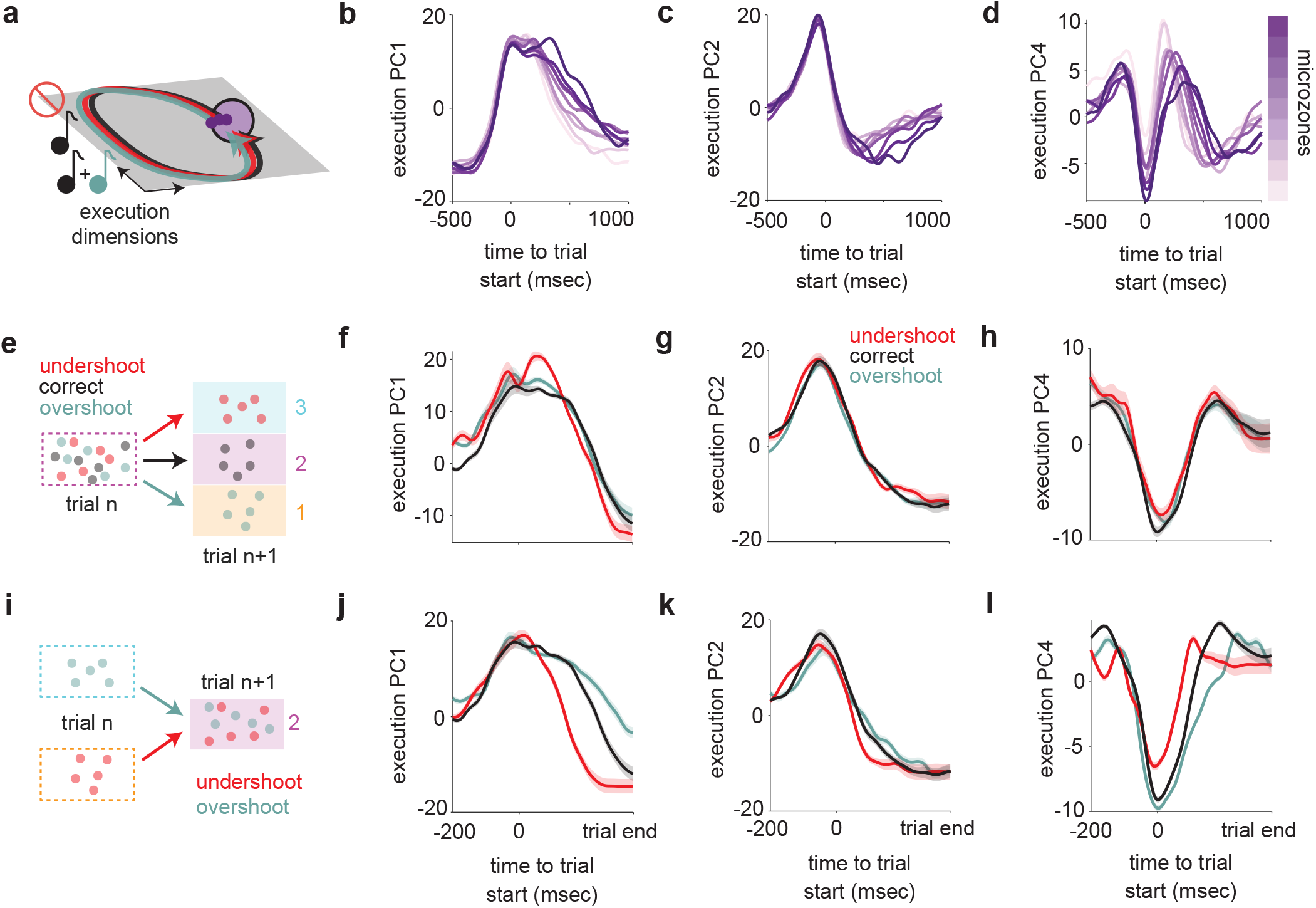
Acoustic errors do not modulate execution dimensions of M2 activity. **a**, Schematic depicting model in which acoustic errors have no effect on motor execution dimensions in 2D space. Trajectories should remain similar for the duration of a trial, unless they diverge due to slight differences in movement durations. **b**, Projection (a.u.) (mean across test/train splits) of microzone specific neural activity onto top principal component derived from trial aligned activity (−500 ms to +1000 ms surrounding trial initiation). **c**, Same as in (**b**), but projections shown for 2nd principal component. **d**, Same as in (**b**) and (**c**), but projections shown for 4th principal component. **e**, Schematic depicting identification of trials landing in the range of zone 2 but intended for different targets. **f**, Projection of M2 activity (a.u.) onto trial aligned PC1 depicted in (**b**) during various trial outcome types requiring correction away from zone 2 range. Shaded regions represent 95% confidence interval across bootstraps (see Methods). Trial durations normalized using the binning procedure described in Methods. **g**, Same as in (**f**), but projections shown for PC2 depicted in (**c**). **h**, Same as in (**f**) and (**g**), but projections shown for PC4 depicted in (**d**). **i**, Schematic depicting identification of trials landing in the range of zone 1 or 3 but intended for zone 2. **j**, Projection of M2 activity (a.u.) surrounding time points of feedback onto PC1 during various trial outcome types requiring correction towards zone 2 range. Shaded regions represent 95% confidence interval across bootstraps. **k**, Same as in (**j**), but projections shown for PC2. **l**, Same as in (**j**) and (**k**), but projections shown for PC4.

**Extended Data Fig. 9.**
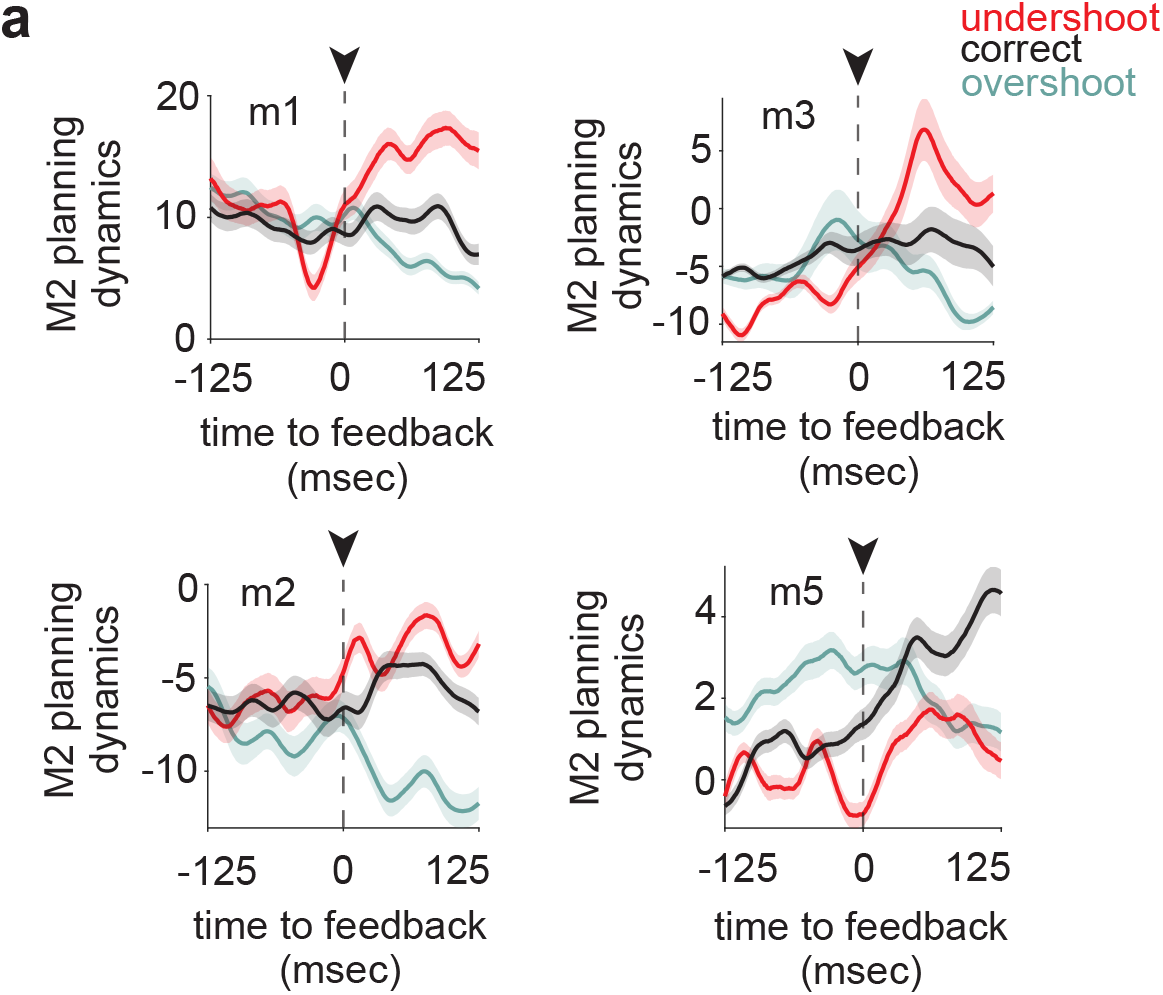
Acoustic feedback interacts with M2 planning dynamics robustly across single animals. **a**, M2 activity (a.u.) projected onto the planning axis (PC1 of pre-trial neural activity), aligned to the moment of acoustic error feedback. All data are from lever presses that apexed within Zone 2, sorted by trial outcome. Shaded regions represent 95% confidence interval across bootstraps (see Methods). Data shown individually for all recordings with at least 10 trials per condition.

**Extended Data Fig. 10.**
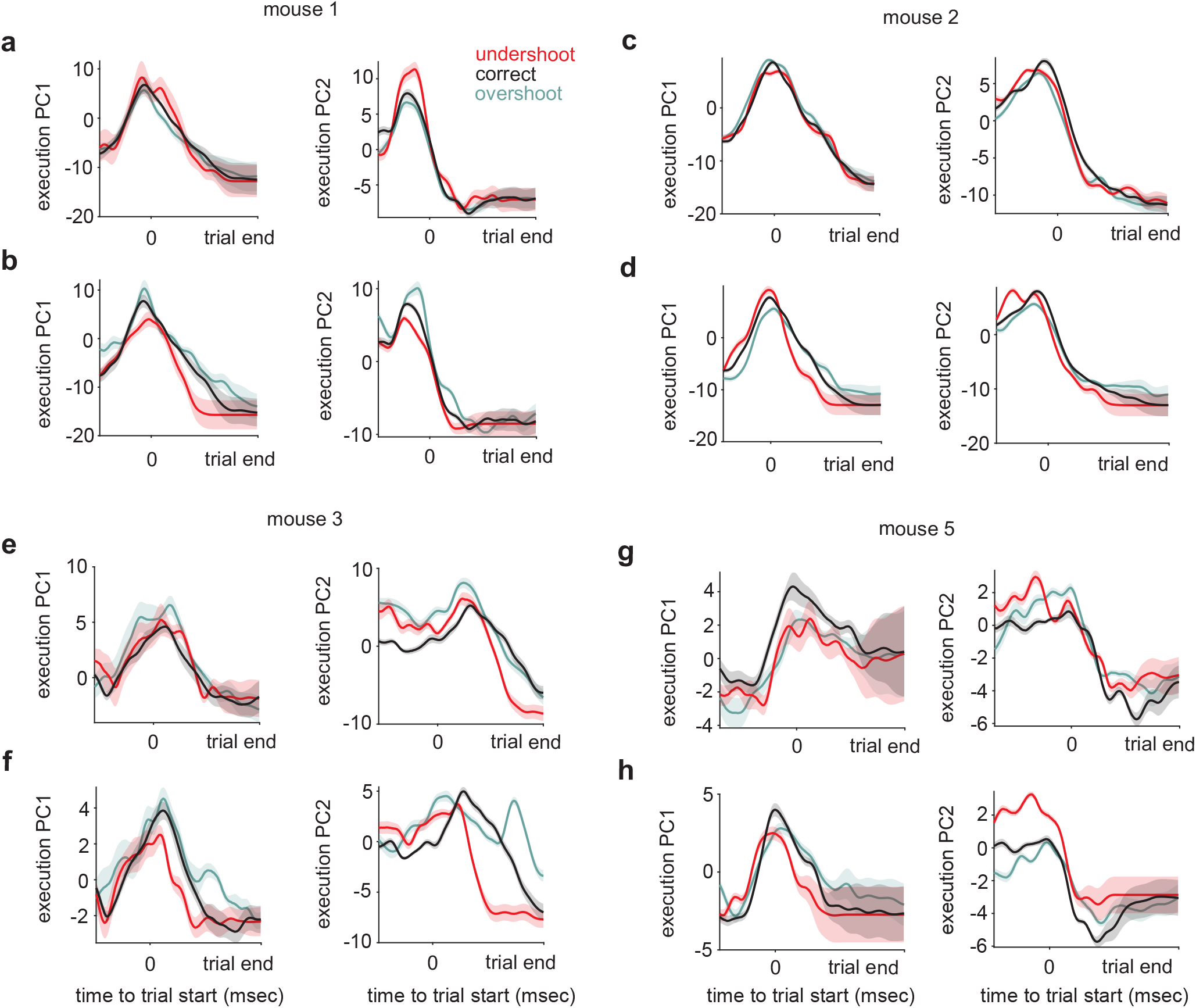
Acoustic errors do not modulate execution dimensions of M2 activity at the single mouse level. **a, c, e, g**, Projection of M2 activity (a.u.) surrounding time points of acoustic feedback onto PC1 & PC2 execution axes (see Extended Data Fig. 8) during various trial outcome types requiring correction away from zone 2 range. Shaded regions represent 95% confidence interval across bootstraps (see Methods). Data shown for all individual recordings with at least 10 trials per condition. Trial durations normalized using ^the binning procedure^ described in Methods. **b, d, f, h**, Projection of M2 activity (a.u.) surrounding time points of feedback onto PC1 & PC2 ^execution^ axes during various trial outcome types requiring correction towards zone 2 range. Shaded regions represent 95% confidence interval across bootstraps (see Methods). Data shown for all recordings with at least 10 trials per condition.

**Extended Data Fig. 11.**
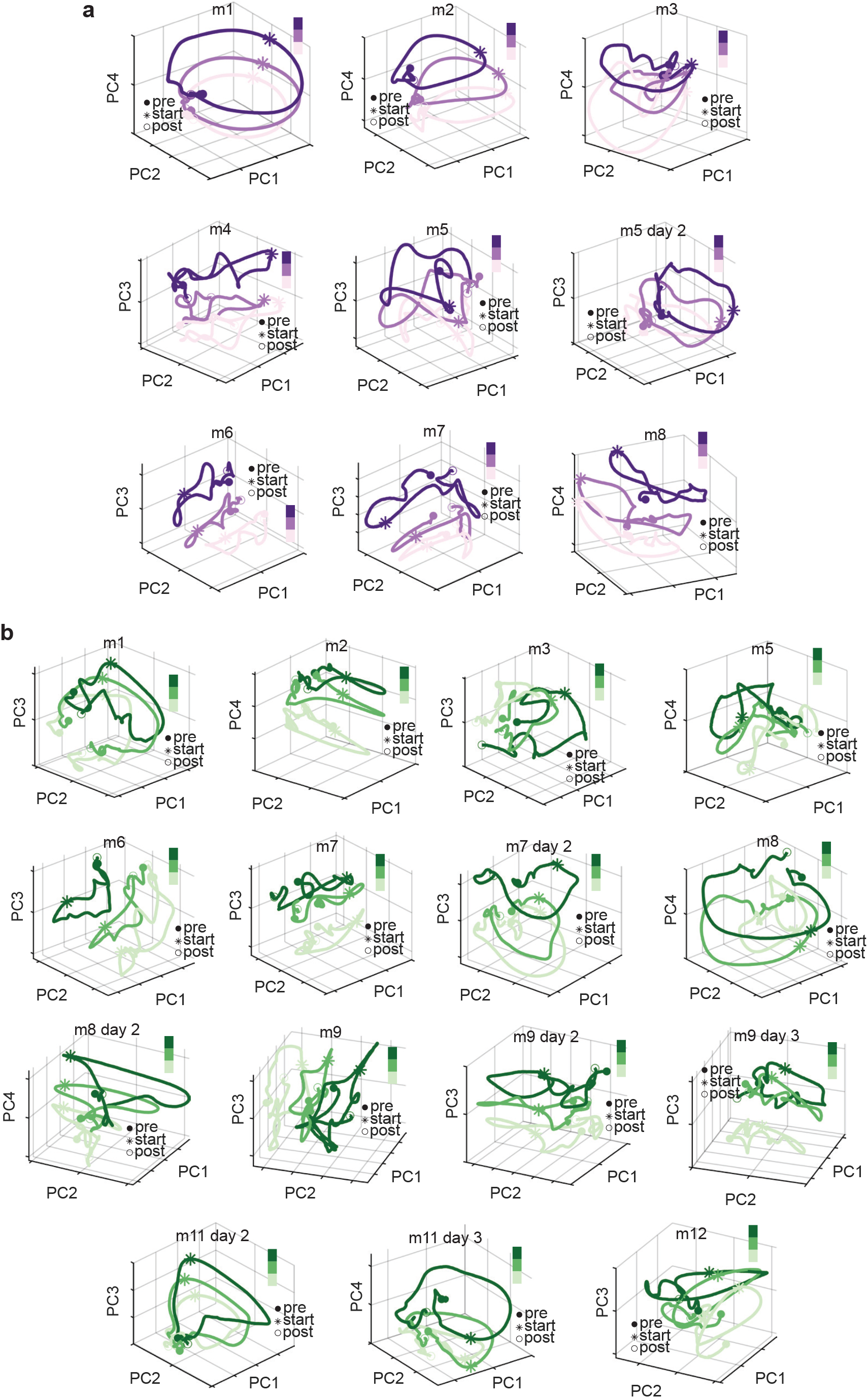
Low-dimensional dynamics and planning-related structure are conserved across individual mice. **a**, Projection of three-zone split neural data into subspace defined by top principal components derived from trial-aligned neural data. Pre (−500 ms), post (+1000 ms) surrounding trial initiation. Data shown for individual mouse (“m”) recordings from secondary motor cortex. **b**, Same as in (**a**), but data shown for individual primary auditory cortex recordings.

**Extended Data Fig. 12.**
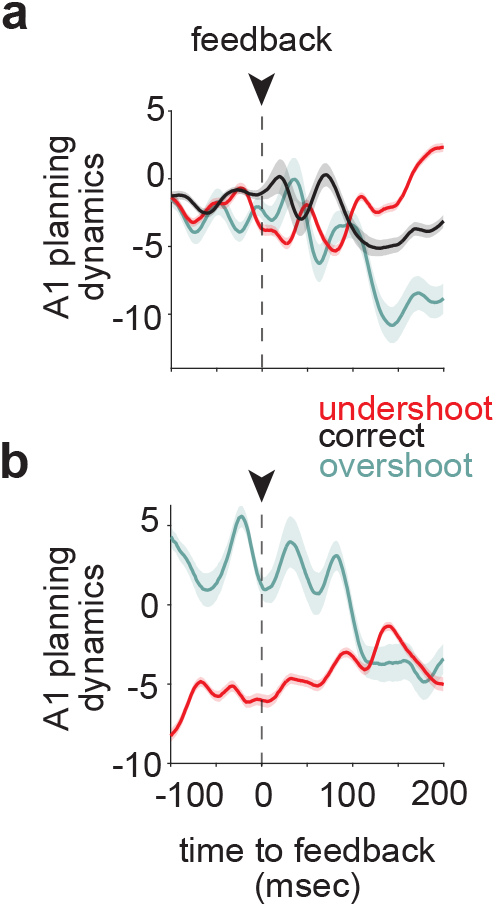
A1 planning dynamics inherit feedback aligned deflections with a delay relative to M2. **a**, Equivalent anaysis to that depicted in **Figure 3n**, but data shown for A1 population activity across all neurons (n=852) with at least 10 trials per condition. A1 population activity is similarly driven by error feedback as M2 is on the planning axis, but with a delay of around 80-100ms. **b**, Same as in (**a**), but data shown for undershoot and overshoot trials that were intended to land in Zone 2.

**Extended Data Fig. 13.**
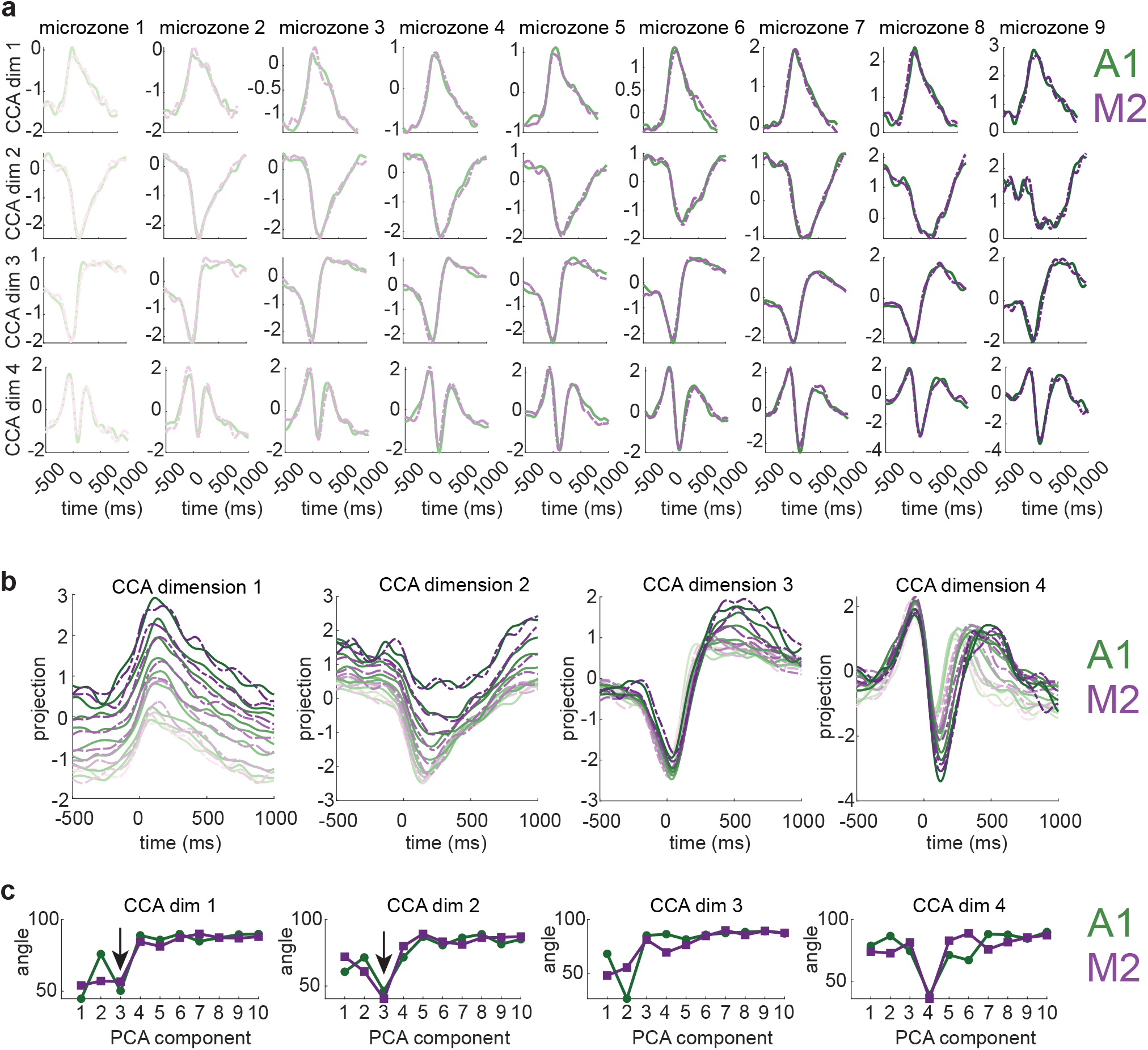
Canonical correlation analysis of A1 and M2 neural activity. **a**, Individual CCA dimension projections (a.u.) across time shown for all 9 microzones, A1 (green) and M2 (purple). **b**, Same as in (**a**), but all microzones plotted on single axis (a.u.). **c**, Angle between individual principal components from each brain region and joint CCA dimensions, A1 (green) and M2 (purple). Arrow indicates similarity between PC3 and CCA dim1 & 2.

**Extended Data Fig. 14.**
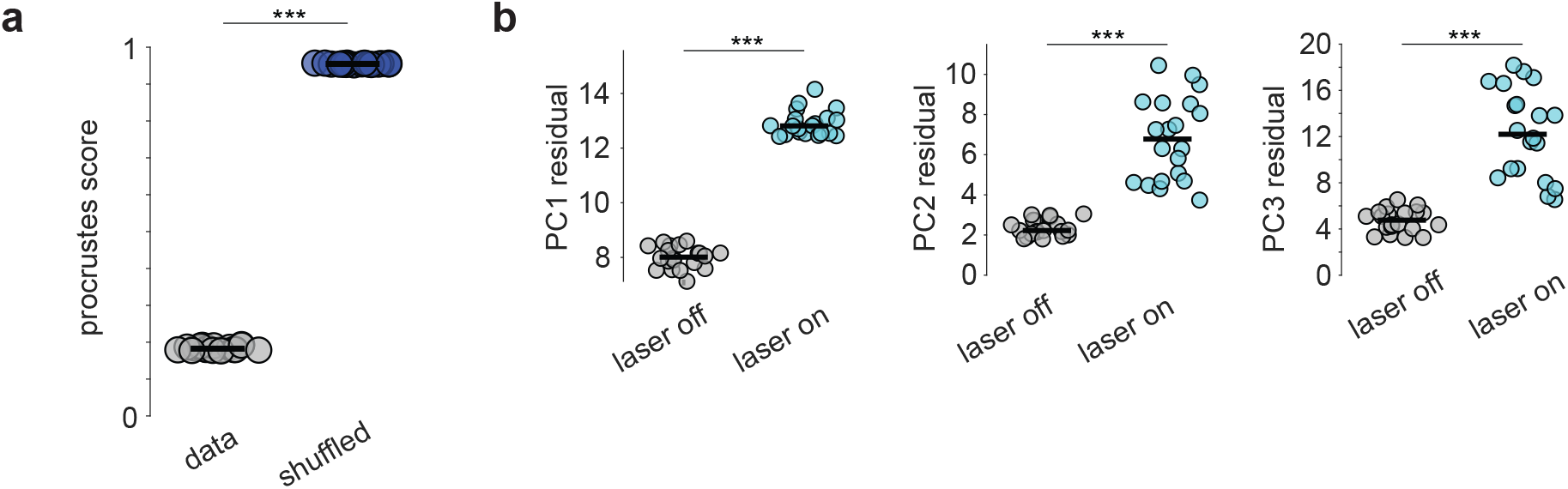
Procrustes alignment of A1-M2 neural manifolds. **a**, Quantification of aligned A1-M2 manifolds vs. shuffled control (see Methods) across 20 data splits, (ranksum, ***p<0.001). Each dot represents alignment metric of neural manifolds trained on a single trial split. **b**, Quantification of individual PC residuals (a.u.) resulting from procrustes alignment of 3-dimensional A1-M2 manifolds under optogenetic silencing conditions depicted in **Figure 4k-n**, comparing laser on/off conditions (ranksum, ***p<0.001). Each dot represents residual resulting from a single data split.

**Extended Data Fig. 15.**
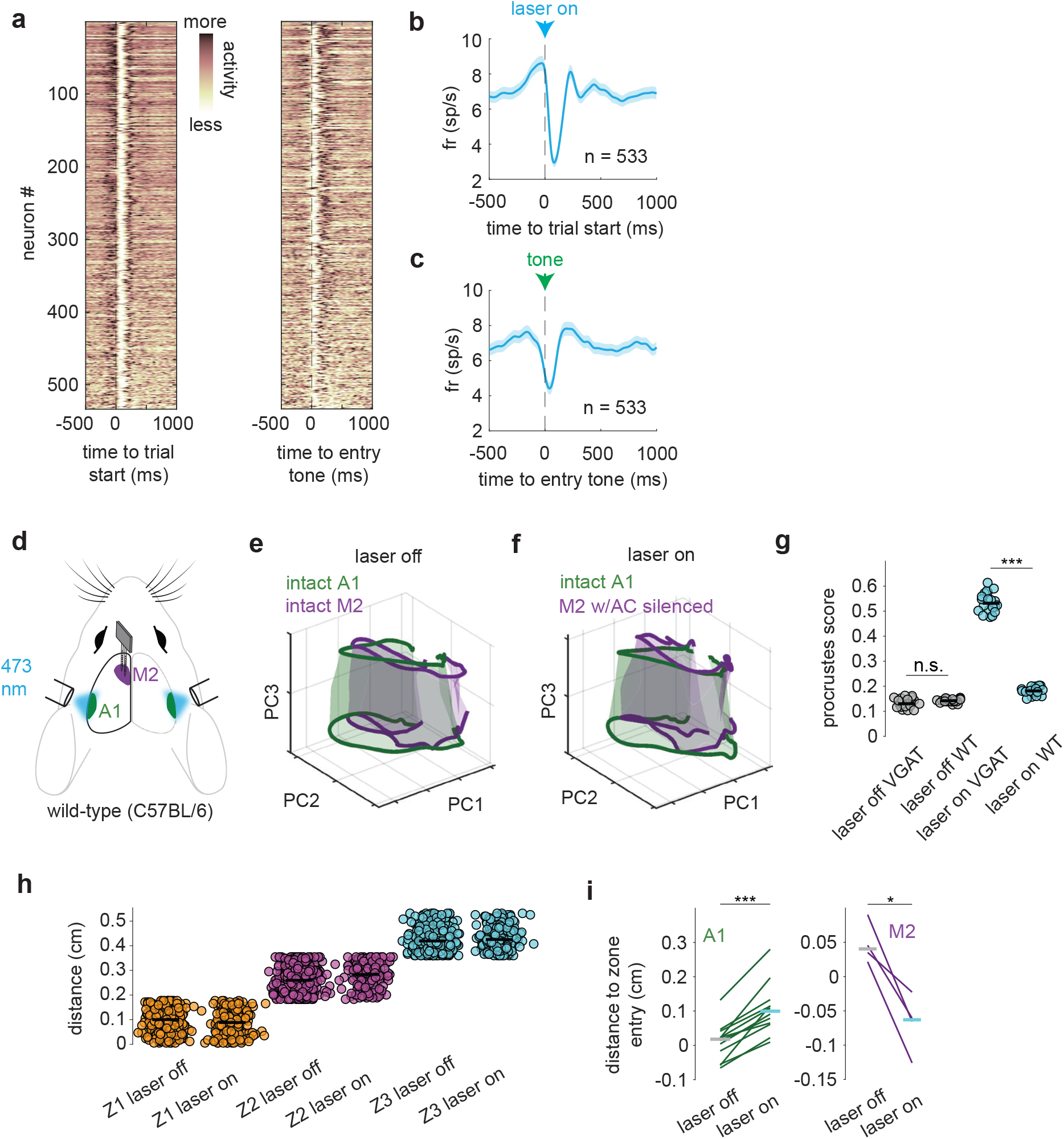
Laser stimulation of auditory cortex in mice not expressing opsins results in no perturbation of the M2 manifold. **a**, Left: M2 population trial averaged PSTHs (n=533) aligned to trial start during AC optogenetic silencing trials. Right: M2 population trial averaged PSTHs (n=533) aligned to time of entry tone on correct trials during AC optogenetic silencing. Each neuron’s PSTH normalized to occupy range of 0-1. **b** and **c**, Population averaged PSTHs for the data shown in (**a**), shaded regions represent SEM. **d**, Schematic depicting bilateral silencing of AC while recording in M2 during behavior in WT/VGAT(−) mice. **e**, Procrustes aligned 3D neural manifolds (0.13 score +/-0.011), intact A1 (green) vs. intact M2 (purple). Reference region: intact A1. Manifolds interpolated across conditions to assist with visualization. **f**, Procrustes aligned 3D neural manifolds (0.17 score +/-0.032), intact A1 (green) vs. M2 with AC silenced (purple). Reference region: intact A1. Manifolds interpolated across conditions to assist with visualization. **g**, Procrustes alignment quantification across 20 data splits, comparing across laser off/on conditions (ranksum, ***p<0.001; n.s. not-significant). Each dot represents alignment metric of neural manifolds in the top 3 PC subspace trained on a single trial split (see Methods). Horizontal bars represent population means. **h**, Peak amplitude of presses included in control vs. optogenetic silencing perturbation experiments in **Figure 4k-n**. Horizontal bars represent population means. **i**, Mean distance of press peak to zone entry position during control and optogenetic perturbation blocks comparing A1 vs. M2 silencing conditions across all mice (paired t-test, ***p<0.001; *p<0.05).

**Extended Data Fig. 16.**
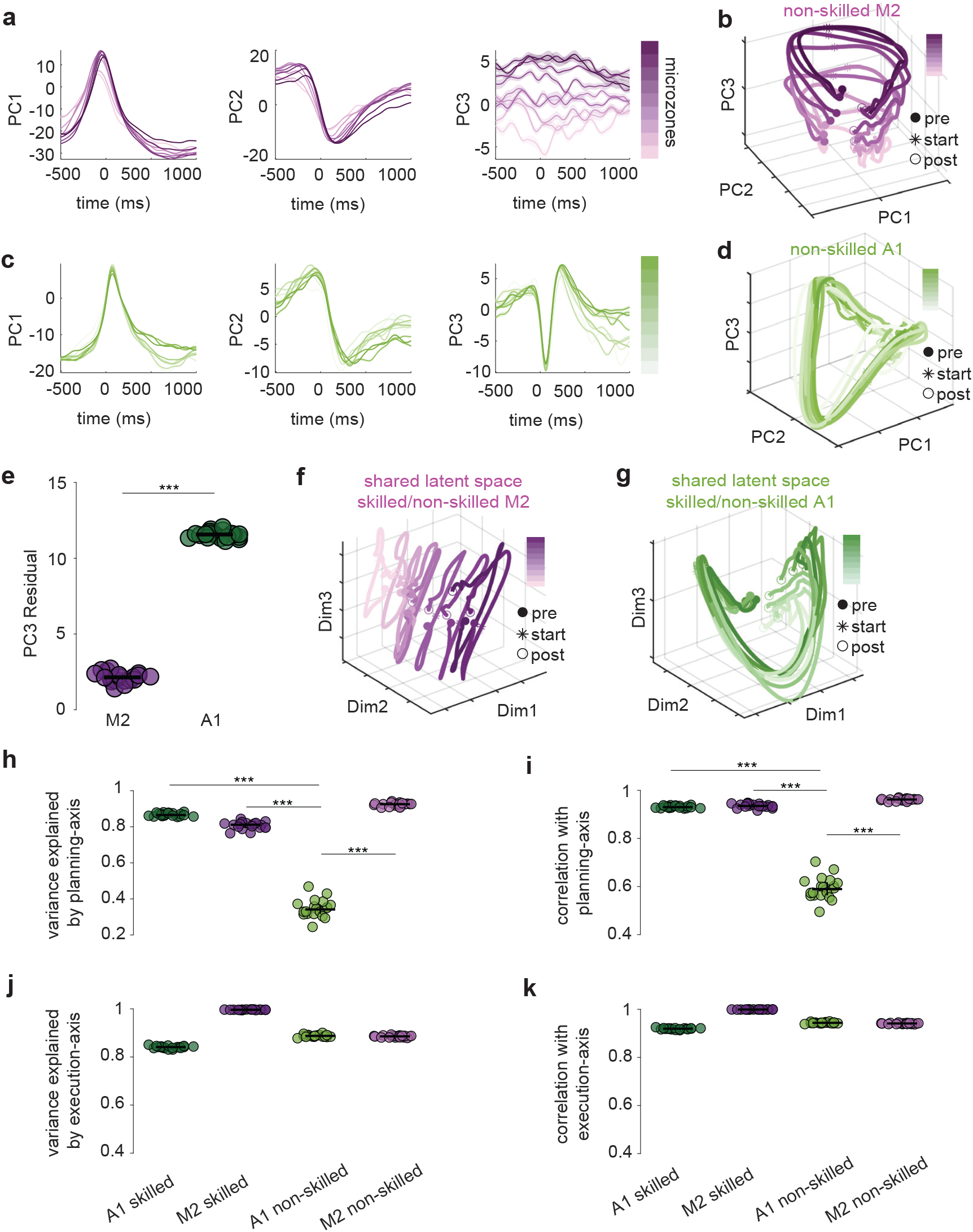
A1 and M2 share manifold geometry only in animals that have learned a skilled, acoustic behavior. **a**, Low-dimensional projections (mean across test/train splits) of non-skilled M2 (n=1326) neural activity onto top 3 principal components individually, 23.8% variance captured by top 3 PCs. **b**, Mean projection of nine-microzone split non-skilled M2 neural data into subspace defined by top 3 principal components. Pre (−500 ms), post (+1000 ms). **c**, Low-dimensional projections (mean across test/train splits) of non-skilled A1 (n=409) neural activity onto top 3 principal components individually, 26.1% variance captured by top 3 PCs. **d**, Mean projection of nine-microzone split non-skilled A1 neural data into subspace defined by top 3 principal components. Pre (−500 ms), post (+1000 ms). **e**, Quantification of PC3 ‘planning-axis’ residual resulting from 3D procrustes alignment of M2 (skilled/non-skilled) manifolds versus A1 (skilled/non-skilled) manifolds, (ranksum, ***p<0.001). **f**, Mean latent space projection into M2 (skilled & non-skilled) joint subspace identified via CCA. Projections for both datasets averaged together and shown as individual condition trajectories. Pre (−500 ms), post (+1000 ms). **g**, Mean latent space projection into A1 (skilled & non-skilled) joint subspace identified via CCA. Projections for both datasets averaged together and shown as individual condition trajectories. Pre (−500 ms), post (+1000 ms). **h**, Fraction of variance in the reference planning-axis projection captured by the Partial least-squares predicted projection across various brain regions/conditions (see Methods), (ranksum, ***p<0.001). **i**, Pearson correlation between the Partial least-squares predicted projection and the reference planning-axis projection across various brain regions/conditions, quantifying the strength and direction of linear correspondence, (ranksum, ***p<0.001). **j**, Fraction of variance in the reference execution-axis projection captured by the Partial least-squares predicted projection across various brain regions/conditions (see Methods). **k**, Pearson correlation between the Partial least-squares predicted projection and the reference execution-axis projection across various brain regions/conditions, quantifying the strength and direction of linear correspondence.

## Bibliography

1. Doupe, A. J. & Kuhl, P. K. Birdsong and human speech: common themes and mechanisms. Annu. Rev. Neurosci. 22, 567–631 (1999).

2. Brainard, M. & Doupe, A. Auditory feedback in learning and maintenance of vocal behaviour. Nat. Rev. Neurosci. 1, 31–40 (2000).

3. Gates, A. & Bradshaw, J. L. Effects of auditory feedback on a musical performance task. Perception & Psychophysics (1974) doi:10.3758/BF03203260.

4. Hubbard, T. L. Auditory imagery: empirical findings. Psychol. Bull. 136, 302–329 (2010).

5. Godøy, R. I. Gestural imagery in the service of musical imagery. in Gesture-Based Communication in Human-Computer Interaction 55–62 (Springer Berlin Heidelberg, Berlin, Heidelberg, 2004).

6. Keller, P. E. Mental imagery in music performance: underlying mechanisms and potential benefits: Keller. Ann. N. Y. Acad. Sci. 1252, 206–213 (2012).

7. Cerkevich, C. M.Rathelot, J.-A. & Strick, P. L. Cortical basis for skilled vocalization. Proc. Natl. Acad. Sci. U. S. A. 119, e2122345119 (2022).

8. Zatorre, R., Chen, J. L. & Penhune, V. When the brain plays music: auditory–motor interactions in music perception and production. Nat. Rev. Neurosci. 8, 547–558 (2007).

9. Nelson, A. & Mooney, R. The basal forebrain and motor cortex provide convergent yet distinct movement-related inputs to the auditory cortex. Neuron 90, 635–648 (2016).

10. Nelson, A. et al. A circuit for motor cortical modulation of auditory cortical activity. J. Neurosci. 33, 14342–14353 (2013).

11. Clayton, K. K. et al. Auditory corticothalamic neurons are recruited by motor preparatory inputs. Curr. Biol. 31, 310–321.e5 (2021).

12. Holey, B. E. & Schneider, D. M. Sensation and expectation are embedded in mouse motor cortical activity. Cell Rep. 43, 114396 (2024).

13. Tian, X. & Poeppel, D. Mental imagery of speech and movement implicates the dynamics of internal forward models. Front. Psychol. 1, 166 (2010).

14. Blakemore, S. J., Wolpert, D. M. & Frith, C. D. Central cancellation of self-produced tickle sensation. Nat. Neurosci. 1, 635–640 (1998).

15. Schneider, D. M. & Mooney, R. Motor-related signals in the auditory system for listening and learning. Curr. Opin. Neurobiol. 33, 78–84 (2015).

16. Keller, G. B. & Mrsic-Flogel, T. D. Predictive Processing: A Canonical Cortical Computation. Neuron 100, 424–435 (2018).

17. Reznik, D., Guttman, N., Buaron, B., Zion-Golumbic, E. & Mukamel, R. Action-locked neural responses in auditory cortex to self-generated sounds. Cereb. Cortex 31, 5560–5569 (2021).

18. Schneider, D. M., Nelson, A. & Mooney, R. A synaptic and circuit basis for corollary discharge in the auditory cortex. Nature 513, 189–194 (2014).

19. Eliades, S. J. & Wang, X. Neural substrates of vocalization feedback monitoring in primate auditory cortex. Nature 453, 1102–1106 (2008).

20. Eliades, S. J. & Wang, X. Dynamics of auditory-vocal interaction in monkey auditory cortex. Cereb. Cortex 15, 1510–1523 (2005).

21. Schneider, D. M., Sundararajan, J. & Mooney, R. A cortical filter that learns to suppress the acoustic consequences of movement. Nature 561, 391–395 (2018).

22. Audette, N. J. & Schneider, D. M. Stimulus-specific prediction error neurons in mouse auditory cortex. J. Neurosci. 43, 7119–7129 (2023).

23. Audette, N. J., Zhou, W., La Chioma, A. & Schneider, D. M. Precise movement-based predictions in the mouse auditory cortex. Curr. Biol. 32, 4925–4940.e6 (2022).

24. Flinker, A. et al. Single-trial speech suppression of auditory cortex activity in humans. J. Neurosci. 30, 16643–16650 (2010).

25. Behroozmand, R. & Larson, C. R. Error-dependent modulation of speech-induced auditory suppression for pitch-shifted voice feedback. BMC Neurosci. 12, 54 (2011).

26. Sober, S. J. & Brainard, M. S. Adult birdsong is actively maintained by error correction. Nat. Neurosci. 12, 927–931 (2009).

27. Alemi, R., Lehmann, A. & Deroche, M. Adaptation to pitch-altered feedback is independent of one’s own voice pitch sensitivity. Sci. Rep. 10, (2020).

28. Khilkevich, A. et al. Brain-wide dynamics linking sensation to action during decision-making. Nature 634, 890–900 (2024).

29. Michaels, J.A., Kashefi, M., Zheng, J. et al. Sensory expectations shape neural population dynamics in motor circuits. Nature 648, 668–677 (2025).

30. Zhou, W. & Schneider, D. M. Learning within a sensory-motor circuit links action to expected outcome. bioRxivorg (2024) doi:10.1101/2024.02.08.579532.

31. Zhou, M. et al. Scaling down of balanced excitation and inhibition by active behavioral states in auditory cortex. Nat. Neurosci. 17, 841–850 (2014).

32. Ceballo, S., Piwkowska, Z., Bourg, J., Daret, A. & Bathellier, B. Targeted cortical manipulation of auditory perception. Neuron 104, 1168–1179.e5 (2019).

33. Letzkus, J. J. et al. A disinhibitory microcircuit for associative fear learning in the auditory cortex. Nature 480, 331–335 (2011).

34. van Sonderen, J. F., Gielen, C. C. & Denier van der Gon, J. J. Motor programmes for goal-directed movements are continuously adjusted according to changes in target location. Exp. Brain Res. 78, 139–146 (1989).

35. Grell, A., Sundberg, J., Ternström, S., Ptok, M. & Altenmüller, E. Rapid pitch correction in choir singers. J. Acoust. Soc. Am. 126, 407–413 (2009).

36. Znamenskiy, P. & Zador, A. M. Corticostriatal neurons in auditory cortex drive decisions during auditory discrimination. Nature 497, 482–485 (2013).

37. Li, N., Chen, T.-W., Guo, Z. V., Gerfen, C. & Svoboda, K. A motor cortex circuit for motor planning and movement. Nature 519, 51–56 (2015).

38. Heindorf, M., Arber, S. & Keller, G. B. Mouse motor cortex coordinates the behavioral response to unpredicted sensory feedback. Neuron 99, 1040–1054.e5 (2018).

39. Barthas, F. & Kwan, A. C. Secondary motor cortex: Where ‘sensory’ meets ‘motor’ in the rodent frontal cortex. Trends Neurosci. 40, 181–193 (2017).

40. Sauerbrei, B. A. et al. Cortical pattern generation during dexterous movement is input-driven. Nature 577, 386–391 (2019).

41. Churchland, M. M. & Shenoy, K. V. Preparatory activity and the expansive null-space. Nat. Rev. Neurosci. 25, 213–236 (2024).

42. Kaufman, M. T., Churchland, M. M. & Ryu, S. I. Cortical activity in the null space: permitting preparation without movement. Nat. Neurosci. (2014).

43. Churchland, M. et al. Neural population dynamics during reaching. Nature 487, 51–56 (2012).

44. Gallego, J. A., Perich, M. G., Miller, L. E. & Solla, S. A. Neural manifolds for the control of movement. Neuron 94, 978–984 (2017).

45. Mitelut, C. et al. Mesoscale cortex-wide neural dynamics predict self-initiated actions in mice several seconds prior to movement. Elife 11, e76506 (2022).

46. Novembre, G. & Keller, P. E. A conceptual review on action-perception coupling in the musicians’ brain: what is it good for? Front. Hum. Neurosci. 8, 603 (2014).

47. Duncker, L. & Sahani, M. Dynamics on the manifold: Identifying computational dynamical activity from neural population recordings. Curr. Opin. Neurobiol. 70, 163–170 (2021).

48. Javadzadeh, M. & Hofer, S. B. Dynamic causal communication channels between neocortical areas. Neuron 110, 2470–2483.e7 (2022).

49. Semedo, J. D., Zandvakili, A., Machens, C. K., Yu, B. M. & Kohn, A. Cortical areas interact through a communication subspace. Neuron 102, 249–259.e4 (2019).

50. Perich, M. G., Narain, D. & Gallego, J. A. A neural manifold view of the brain. Nat. Neurosci. (2025) doi:10.1038/s41593-025-02031-z.

51. MacDowell, C. J. et al. Multiplexed subspaces route neural activity across brain-wide networks. Nat. Commun. 16, 3359 (2025).

52. Petersen, C. C. H. Sensorimotor processing in the rodent barrel cortex. Nat. Rev. Neurosci. 20, 533–546 (2019).

53. Cohen, M. X. & Ranganath, C. Reinforcement learning signals predict future decisions. Journal of Neuroscience (2007).

54. Dryden, I. L. & Mardia, K. V. tStatistical shape analysis: with applications in R. (2016).

55. Driscoll, L. N., Shenoy, K. & Sussillo, D. Flexible multitask computation in recurrent networks utilizes shared dynamical motifs. Nat. Neurosci. 27, 1349–1363 (2024).

56. Drieu, C. et al. Rapid emergence of latent knowledge in the sensory cortex drives learning. Nature 641, 960–970 (2025).

57. McGinley, M. J. et al. Waking state: Rapid variations modulate neural and behavioral responses. Neuron 87, 1143–1161 (2015).

58. Otazu, G. H., Tai, L.-H., Yang, Y. & Zador, A. M. Engaging in an auditory task suppresses responses in auditory cortex. Nat. Neurosci. 12, 646–654 (2009).

59. Svoboda, K. & Li, N. Neural mechanisms of movement planning: motor cortex and beyond. Curr. Opin. Neurobiol. 49, 33–41 (2018).

60. Inagaki, H. K., Fontolan, L., Romani, S. & Svoboda, K. Discrete attractor dynamics underlies persistent activity in the frontal cortex. Nature 566, 212–217 (2019).

61. Elsayed, G. F., Lara, A. H., Kaufman, M. T., Churchland, M. M. & Cunningham, J. P. Reorganization between preparatory and movement population responses in motor cortex. Nat. Commun. 7, 13239 (2016).

62. Shepherd, G. & Yamawaki, N. Untangling the cortico-thalamo-cortical loop: cellular pieces of a knotty circuit puzzle. Nat. Rev. Neurosci. 22, 389–406 (2021).

63. Prather, J., Peters, S., Nowicki, S. & Mooney, R. Precise auditory–vocal mirroring in neurons for learned vocal communication. Nature 451, 305–310 (2008).

64. Rao, R. P. N. A sensory-motor theory of the neocortex. Nat. Neurosci. (2024) doi:10.1038/s41593-024-01673-9.

65. Ames, K. C., Ryu, S. I. & Shenoy, K. V. Simultaneous motor preparation and execution in a last-moment reach correction task. Nat. Commun. 10, 2718 (2019).

66. Stavisky, S. D., Kao, J. C., Ryu, S. I. & Shenoy, K. V. Motor cortical visuomotor feedback activity is initially isolated from downstream targets in output-null neural state space dimensions. Neuron 95, 195–208.e9 (2017).

67. Pachitariu, M., Sridhar, S., Pennington, J. & Stringer, C. Spike sorting with Kilosort4. Nat. Methods 21, 914–921 (2024).

68. Kalatsky, V. A. & Stryker, M. P. New paradigm for optical imaging: temporally encoded maps of intrinsic signal. Neuron 38, 529–545 (2003).

